# A Computational Pipeline for Physiologically Informed Calibration of Ligand Reaction-Diffusion Models Using High-Throughput Sequencing

**DOI:** 10.1101/2025.08.25.672053

**Authors:** Ali Daher, Dumitru Trucu, Raluca Eftimie

## Abstract

All physiological processes fundamentally rely on continuous cellular cross-talk to maintain organization and ensure proper function. Among the various modes of cellular communication, ligand-mediated chemical signaling, in which a ligand is secreted by one cell, diffuses through the extracellular environment, and binds to a receptor on another (or the same) cell to elicit a downstream response, is arguably the most ubiquitous and foundational. Given its importance, numerous mathematical models have been developed to describe this reaction-diffusion mechanism, capturing ligand secretion, diffusion, decay, and binding under both normal and pathological conditions. However, parameter calibration for such models often lags behind model development. This is due to limited data that faithfully represent the biological microenvironment, as well as due to the absence of a robust, rigorous framework to integrate available data into the mathematical model. To address this gap, we propose that transcriptomics (gene expression) data, namely the combination of single-cell RNA sequencing and spatial transcriptomics, provide a rich, increasingly abundant, and underutilized source of information that can be used to calibrate the parameters of the cellular reaction-diffusion models at the larger mesoscopic scale. To this end, we develop a computational pipeline that leverages these data to extract parameter values for reaction-diffusion models, and illustrate its application through two human wound-healing case studies. Using open-source transcriptomics data, we calibrate the reaction-diffusion model parameters of the isoforms of Transforming Growth Factor Beta (TGF*β*), a signaling molecule central to tissue repair and development as well as to pathological processes such as cancer and fibrosis. Our pipeline integrates traditional numerical (finite volume) solvers for the ligand concentration fields with bioinformatics, machine learning, and Bayesian inference methods, combining existing and novel computational tools into a single framework for a physiologically informed, data-driven parameter calibration process. The pipeline is modular, allowing easy extension or adjustment depending on user needs. Overall, this framework facilitates rigorous model calibration, an essential step toward ensuring that mathematical models have meaningful research and potential translational utility.

## 1 Introduction

Cellular cross-talk plays a fundamental role in coordinating tightly regulated processes of tissue organization and repair. To make decisions, cells communicate through a variety of signalling modalities, depending on the physiological context and spatial range of communication. These include ligand-mediated paracrine and autocrine chemical signalling, contact-based juxtacrine signaling, neuronal synaptic communication, endocrine signalling in the form of hormones travelling through the blood stream and, as more recently unravelled, long-range mechanical traction forces through supracellular filaments that influence collective cell migration.^1^ Among these signalling pathways, paracrine and autocrine chemical signalling are particularly foundational within tissue microenvironments, mediating local interactions between neighbouring cells (paracrine) or within the same cell (autocrine). In both of these modalities, a cell releases a ligand, such as a growth factor, which then diffuses through the extracellular space and binds to a complementary receptor, eliciting a downstream signalling cascade in the target cell. This form of short-range chemical signalling is central to numerous biological processes, including development, inflammation, wound healing, and tissue remodeling. Dysregulations in these pathways have been implicated in a range of pathological conditions, including cancer, fibrosis, and chronic inflammatory diseases. ^2–6^

Given its central role in tissue regulation, chemically mediated cellular cross-talk has been the focus of extensive mathematical and computational modelling efforts.^7–9^ One of the most widely used approaches involves reaction-diffusion models, which characterize the production, diffusion, degradation, and receptor binding of signalling ligands using particle or continuum-based spatial or spatiotemporal partial differential equations (PDEs).^10, 11^ These equations can be used to compute the spatial ligand concentration fields across space and time. In addition, they can also be coupled to cellular dynamics equations capturing cellular proliferation, migration, and death, in order to incorporate the effects of chemical signalling on these processes.

For such models to be clinically or scientifically meaningful, their parameter values must be carefully calibrated in a physiologically relevant and data-informed manner. Historically, however, the parameter selection process in many of these mechanistic models has often lacked behind model development in terms of rigour, robustness, and biological fidelity.^12^ Traditionally, parameter values have been calibrated using one or both of the following approaches. First, they are sometimes heuristically tuned through trial and error in computational simulations; for instance, by selecting values that avoid numerical instability or that produce model outputs consistent with broad biological expectations. Yet, the true strength of in silico models lies in their ability to probe biological mechanisms whose outputs are unknown or poorly understood; thus, reliance on expected behaviour limits their exploratory power.

Second, in some cases, parameter values are indeed derived from in vitro experimental assays, such as ligand-receptor binding studies used to estimate binding kinetics; see^10, 13^ for examples. However, this approach presents its own limitations. In vitro conditions fail to fully recapitulate the complexity of the tissue biological environment. Moreover, these experiments are often conducted under highly specific conditions, varying by species, cell type, or tissue context, making it difficult to generalize the resulting parameters values. In many cases, experimental data are only available for a subset of the required parameters, leaving the remainder uncertain. Even when data are available, reported values can span several orders of magnitude depending on the experimental design. For example, the binding rates for transforming growth factor Beta (*TGFβ*) isoform *TGFβ*1 to the *TGFβ* type II receptor (*TGFβ*1 −*R*2) have been reported to range from 5.1×10^5^ to 3.1×10^7^ (*Ms*)^*−*1^, depending on whether or not the receptors in experimental binding assay were immobilized.^13–17^

The gap between model development and parameter calibration raises an important question: what alternative sources of data can be leveraged to tune those parameter values, and what methodological framework can be employed to integrate such data into the model for robust parameter inference? In this paper, we demonstrate how high-throughput genetic sequencing data, specifically the combination of single-cell RNA sequencing (scRNA-seq) and Visium spatial transcriptomics (ST), can be leveraged to calibrate the parameters of reaction-diffusion-based models of chemical signalling between cells.

To this end, we develop a computational pipeline that integrates existing tools with newly devised algorithms in order to bridge the gap between single-cell gene expression data and higher-scale model dynamics, thereby enabling robust parameter calibration. In Section 2, we provide a detailed description of the sequential steps in the pipeline, along with a step-by-step case study that demonstrates how high-throughput genetic data from a skin tissue sample undergoing normal wound healing and collected at day 30 post-incision^18^ can be leveraged to calibrate the parameters of the reaction-diffusion model during the wound remodelling phase. For the case-study, we focus on the three human isoforms for transforming growth factor Beta (TGF*β*), a key regulatory cytokine that plays an essential role across all three stages of the skin wound healing (inflammation, proliferation, and remodelling) and homeostasis, as well as mediates a diverse series of cellular responses including growth suppression, changes in cell morphology, and expression of cell adhesion genes,^19^ making it a crucial regulator of fibrosis, cancer progression, and immune modulation.

Finally, in Section 3, we discuss certain aspects of the computational pipeline, highlight its advantages as well as some of its limitations, and explore potential future work. Overall, our results demonstrate that high throughput genetic sequencing data offer a rich, underutilized and nowadays increasingly accessible (and often freely available) source of information in modern biology for parameter calibration. All the high throughput datasets in this study were obtained from open-source databases, underscoring the broad accessibility of this approach. The computational pipeline developed draws on methods from bioinformatics, machine learning, numerical analysis, and Bayesian inference to facilitate a rigorous and robust parameter calibration process that is physiologically informed and data-driven. While showcased here in the context of skin (normal and abnormal) wound healing, the pipeline is broadly applicable to other biological systems characterized by complex, multicellular dynamics, such as cancer growth and tissue regeneration.

## 2 The Computational Pipeline

The first stage of the pipeline, detailed in Section 2.1, employs conventional bioinformatics methods and existing tools to extract two key data components from the sequencing scRNA and ST datasets. First, it computes the spatial cell density distributions of the relevant cell types across the tissue domain. Second, it estimates the ligand-mediated interaction strengths between different cell types and ligand–receptor pairs of interest, which would then serve as the experimentally derived reference data against which the model outputs can be quantitatively calibrated.

In Section 2.2, we formulate a finite volume (FV) scheme to compute the ligand concentration fields. The spatial discretization (mesh) of the FV solver mirrors the hexagonal grid used in the Visium ST experimental platform, which enables direct comparison and data integration between the model outputs and the experimental data. For each multi-dimensional realization in the parameter space, the FV scheme is used to solve for the spatial concentration field of the growth factor(s) of interest. The input parameter realization, along with the numerically computed growth factor concentration fields, can be used to calculate the interaction strengths across all the relevant cell type pairs and ligand-receptor combinations of interest; these, in turn, can be directly compared to the experimentally derived interaction strengths in Section 2.1.

Next, in Section 2.3, we conduct a thorough review of the existing literature to extract the relevant parameter values of the reaction-diffusion model by examining both experimental and computational studies. These values are used as a starting point to construct an informed prior distribution of the model parameters, from which an initial ensemble of realizations can be generated and subsequently updated following the calibration process.

All that remains is a framework through which the model-derived and the transcriptomics data-derived interaction strengths can be directly compared, with their differences systematically minimized and mapped back to the underlying model parameters. In Section 2.4, we discuss three relevant parameter calibration strategies: the rejection sampler scheme, the Sequential Monte Carlo scheme (SMC), and the gradient descent. The first two fall under a class of computational Bayesian inference techniques known as Approximate Bayesian Computation (ABC), while the latter is a fundamental optimization algorithm for machine learning methods. We outline the advantages and limitations of each method in the context of calibrating parameters in the reaction-diffusion model and propose an integrated calibration pipeline where the three methods are applied sequentially in a hierachial framework, such that the output of one method as the input for the next, in order to harness their individual strengths and mitigate their respective drawbacks.

### 2.1 Bioinformatics Analysis

In this first part, we describe how high-throughput genetic sequencing data, namely, scRNA-seq and ST, can be effectively processed to extract relevant information that ca be used in the subsequent numerical analysis and parameter calibration sections. Specifically, we outline how established bioinformatics methodologies and existing tools can be employed to compute two key quantities: (1) the spatial distribution of cells across the cell types of interest, which is needed to set up the finite volume (FV) numerical scheme, and (2) the interaction strengths for each ligand-receptor and cell-cell pair, approximated from the mRNA co-expression levels of the ligand in the secreting (sending) cells and the corresponding receptor in the receiving cells; these interaction strengths then serve as the reference data against which the model output is compared and calibrated.

#### 2.1.1 ScRNA-Seq Data Processing

In a unique study on human skin wound healing, Landen and colleagues^18^ have induced wounds in healthy donors and collected wound-edge tissues at the three stages: inflammation (day 1), proliferation (day 7), and remodelling (day 30), monitoring the molecular and cellular dynamics of human and skin wound healing at each stage at an unprecedented resolution by performing both ScRNA-seq and ST (Visium) experiments on the samples from two donors at each of the wound-healing stages.^18^ In this study, we focus on analyzing the sample datasets for one of the two donors (donor 3) at day 30 post wound incision, when the wound has already entered the remodelling phase. We choose to analyze the tissue sampe at day 30 because experimental and computational studies^10, 20, 21^ suggest that during this period, the TGF*β* levels and cell population dynamics stabilize and TGF*β* levels approach stable, non-zero concentrations. This means that we can simplify the governing equations and assume quasi-steady state dynamics for the TGF*β* field, governed by a time-invariant reaction-diffusion system (*∂g/∂t* ≈ 0 where *g* is the TGF*β* concentration), while retaining the flexibility to extend the framework to fully time-dependent dynamics as needed.

The goal of the scRNA-seq data processing step is to derive an annotated list, organized by cell type, of gene expression profiles (expressed in gene expression units) for the cells in the tissue sample, starting from the raw, unannotated gene expression matrix. The processing workflow follows largely standardized protocols and aligns with those employed in previous scRNA-seq studies aimed at characterizing the cellular and molecular landscapes of skin tissues and lesions^18, 22, 23^:

1. **Filtering**: Low-quality cells expressing *<* 400 or *>* 8000 genes, ≥20% mitochondrial genes, and *<* 500 gene counts, as well as genes expressed in fewer than 10 cells, were excluded. Potential doublets detected using Scrublet^24^ were also excluded.
2. **Data Normalization and Scaling** were performed using the Scanpy package^25^ in Python.
3. **Dimensionality Reduction and Clustering**: The top 4000 variable genes were used for Principal Component Analysis (PCA) using the Seurat flavour from Scanpy,^25^ and the top 50 components (with the largest eignevalues) were subsequently used in the clustering analysis. Uniform Manifold Approximation and Projection (UMAP) and k-nearest neighbour graph were generated using the Scanpy package. The major clusters were identified using the Louvain igraph-based algorithm in Scanpy, with a resolution of 0.8.
4. **Mapping Clusters to Cell Types**: Differentially expressed genes within each cluster were determined using the Wilcoxon method in Scanpy. To label each cluster by the cell type, the 50 most highly expressed genes in each cluster were determined and mapped against the differential gene expression (DGE) annotation file provided by the original study^18^; these, in turn, were originally obtained by using the ‘FindTransferAnchor’ and ‘MapQuery’ functions in Seurat.^18, 26^
5. The UMAP embedding of the different clusters and their allocated cell type annotation are shown in Figure 1(a), and the cell count per type across the tissue sample is shown in the bar chart in Figure 1(b).

**Figure 1.**
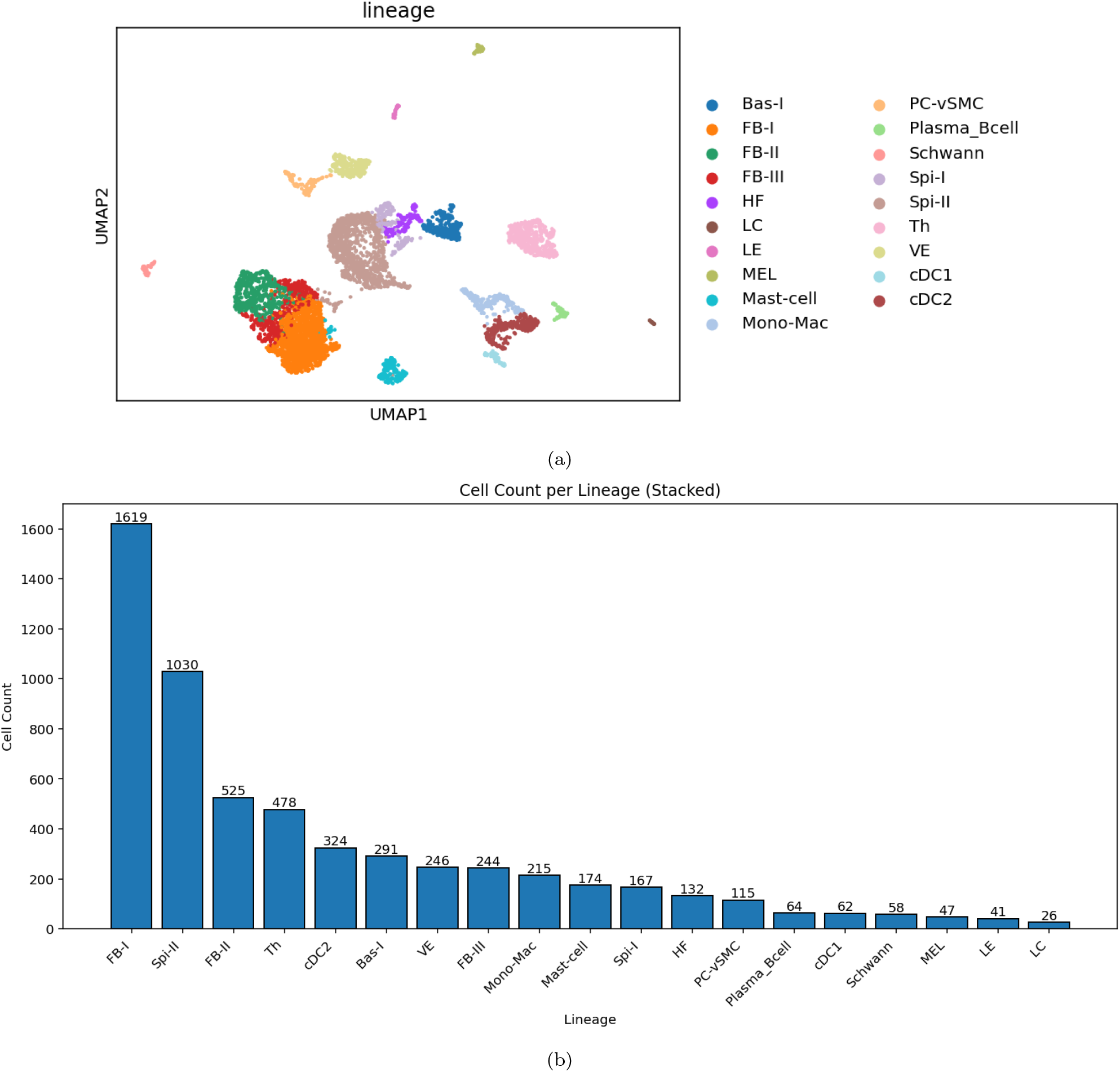
The processed raw scRNA-seq data (for donor 3 at day 30 from skin incision) in the study by Liu et al.^18^ (a) The UMAP embedding of the identified clusters with the corresponding cell type annotations. (b) The cell count per cell type. Bas/Spi: Basal/Spinuous Keratinocytes, FB: Fibroblasts, HF: hair follicle, LC: Langerhan cells, LE/VE: Lymphatic/Vascular Endothelial cell, MEL: Melanocyte, Mono-Mac: Monocytes and Macrophages, PC-vSMC: Pericytes and Vascular Smooth Muscle cells, Th: T-helper cells, cDC: conventional Dendritic Cells.

#### 2.1.2 Reconstructing Spatial Patterns of Cell Densities

Unfortunately, single-cell RNA sequencing (scRNA-seq) data lack spatial context and do not provide information regarding the physical locations or distributions of individual cells within the tissue. To overcome this limitation, many high-throughput transcriptomic studies now integrate scRNA-seq with ST, most commonly using the Visium platform,^27^ which enables spatially resolved gene expression profiling. In the most widely used commercial implementation of Visium, a tissue section is mounted on a glass slide patterned with an array of 55 *µm*-diameter spots arranged in a hexagonal grid. Each spot captures mRNA from nearby cells, allowing the construction of a spots-by-genes expression matrix that retains spatial information. Figure 2(a) shows a histological image of normally healing tissue for donor 30 at day 30 post-incision, while Figure 2(b) displays the corresponding Visium spots arranged in a hexagonal grid and overlayed on the tissue section, illustrating how spatial gene expression is measured across the wound bed.^18^

**Figure 2.**
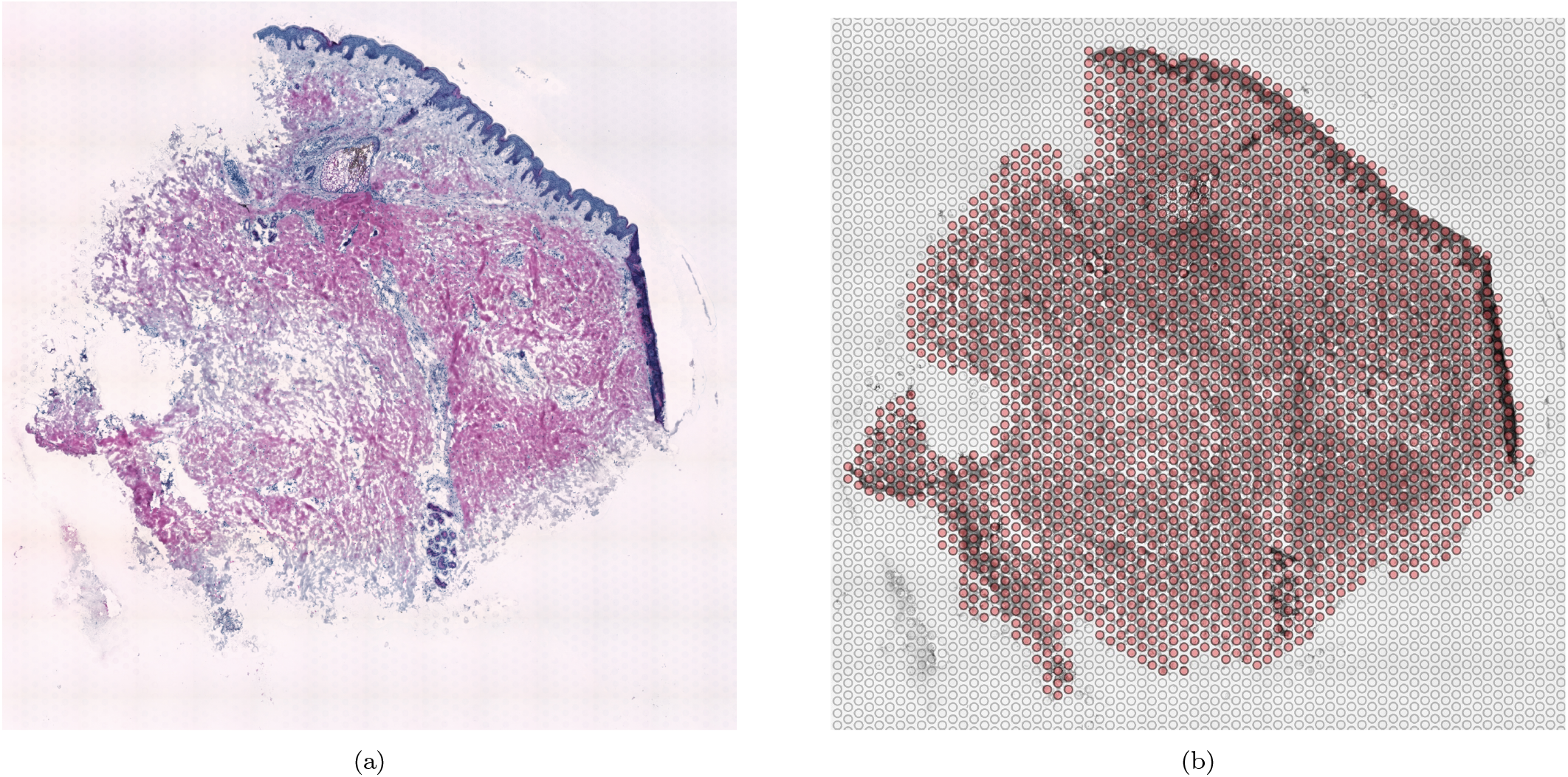
Histological tissue sample representing normal wound healing for donor 3 at day 30 post incision (a), with the corresponding Visium spots in red arranged in a hexagonal grid and overlayed on the tissue sample (b). Figures derived from GEO accession GSE241132, originally published by Liu et al.^18^

To map the spatial positions of the identified cells (according to the scRNA-seq analysis in Section 2.1.1) in their native environments (*i*.*e*., the histological tissue sample in Figure 2(a)), we use a computational toolkit called Cell2location.^28^ Cell2location employs a Bayesian hierarchial model that decomposes the spatial transcriptomics data into contributions from each of the identified cell types, estimating the proportions (or, in case prior information about the cell numbers is provided, the absolute numbers) of cells for each cell type at each Visium spot. In brief, Cell2location uses the top 100 marker genes for each cell type, obtained from the annotated (by cell type) cell-gene count matrix constructed in Section2.1.1, to train a negative binomial regression model and estimate reference transcriptomic signatures. These reference signatures are then used to decompose the observed mRNA counts in each Visium spot and infer the abundance of each cell type.^28^ The model takes the following as input: 1)the raw [*spots×genes*] ST gene count matrix, 2)the annotated [*cells×genes*] scRNA-seq gene count matrix that was laballed from Section 2.1.1, and 3)a prior distribution of the number of cells per spot (either uniform or spot-specific). The output is a posterior distribution for the number of cells of each type in each Visium spot in the domain.

In our analysis, all parameters were set to default except for the following: 1) the expected cell abundance per location, which was set to 20, in line with the approximate average number of nuclei per spot detected in the H&E images in the original experimental study,^18^ and 2) the per-location normalization regularization parameter (*α*_detection_ = 20), which was also adjusted in the original study to account for large variations in mRNA detection sensitivity across different Visium spots.^18^ Furthermore, in this study, we will use the 5% quantile cell number of the posterior distribution to summarize the spatial distribution of cell types across the Visium spots. This choice reflects a conservative estimate that emphasizes high-confidence cell presence^18^ while mitigating the influence of outliers or uncertain assignments. Nonetheless, our computational pipeline can readily accommodate the use of the full posterior distribution of the cell numbers with some corresponding adjustments if desired. The spatial distributions for the selected cell types (fibroblasts, macrophages, and endothelial cells) within the tissue sample are shown in Figure 3. The cell densities (per sample thickness) can be obtained from the cell counts by dividing the latter by the spot area.

**Figure 3.**
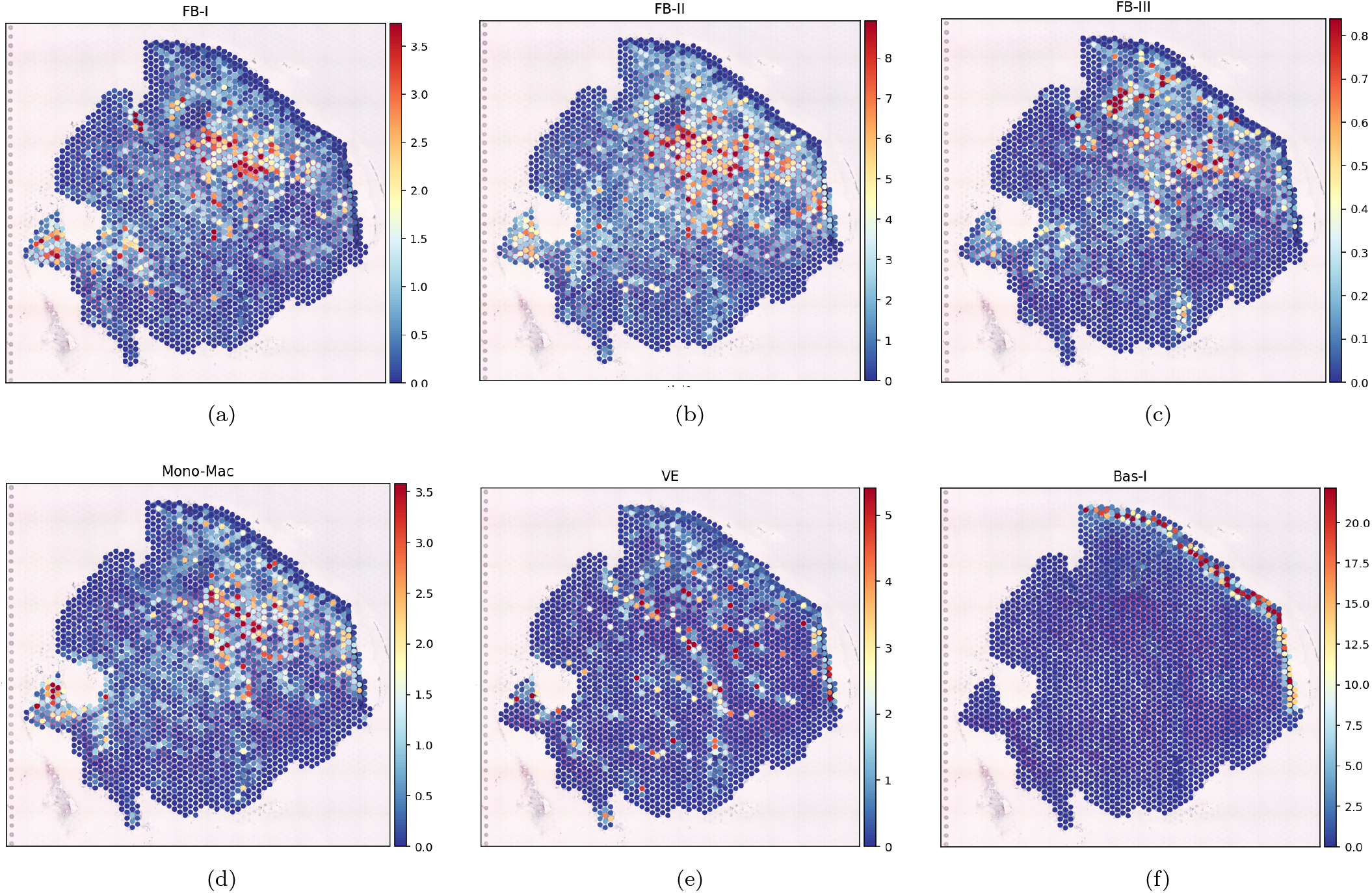
The spatial distribution across the Visium spots of six cell types in the normal-healing tissue sample (donor 3 at day 30 post wound-incision).^18^ The cell counts represent the 5% quantile of the posterior distribution obtained from the Cell2location analysis.

#### 2.1.3 Inferring the Interaction Strengths from the scRNA-seq data

To infer the *TGFβ*-mediated cell-cell interaction strengths from the transcriptomics data, we use CellphoneDB,^29^ a pioneering and widely used computational toolkit for reconstructing cell-cell communication networks across the different cell types from scRNA-seq data. The toolkit leverages a comprehensive curated and annotated database of ligands and receptors interactions, including multi-subunit heteromeric complexes, that is complied from public literature resources, to carry out a systemic analysis of cell-cell communication as motivated by the secreted diffusing ligands.^30^ The ligand-receptor interaction strengths between one secreting cell type and another receiving cell type are derived on the basis of the mean mRNA co-expression of the receptor (or receptor subunits) in one cell type and the ligand in another; see ^31^ for details on the specific calculation of interaction strengths. Note that these interactions are directional and not necessarily symmetric (*i*.*e*., interaction strength from cell type A to B is not the same as from B to A).

In our case, the CellPhonedb algorithm would take the following as input: 1) the annotated (by cell type) cell-gene expression matrix obtained from processing the scRNA-seq data (in Section 2.1.1), 2) The DGE file that features the genetic signature for each cell type, which was used for clustering annotation in Section 2.1.1, and 3) an annotated database of the relevant ligand-receptor interactions, for which we use the default CellphoneDB database. To ensure biological relevance, only ligand–receptor pairs in which both components are expressed in more than 10% of cells within their respective clusters are considered. Furthermore, at least one component of the ligand–receptor pair must be listed in the DGE file for the interaction to be considered.

CellPhoneDB uses the co-expression of ligand–receptor pairs between cell types as a proxy for estimating interaction strength. While this approach provides a practical and interpretable measure and has its advantages, it also comes with limitations, which are discussed in Section 3, along with other potential toolkits that can be used. Nonetheless, at this stage, we chose CellPhoneDB due to its widespread adoption, ease of integration, and suitability for proof-of-concept demonstrations of our computational pipeline.^6, 23, 29, 30^

In Appendix A, we further illustrate the utility of our computational pipeline using another open-source high-throughput sequencing dataset, derived from a skin tissue sample of Acne keloidalis, a chronic inflammatory condition primarily affecting the scalp and neck, characterized by papules, pustules, hair loss, and keloidal scarring.^23^ In this example, interaction strengths are spatially inferred using a recently developed bioinformatics tool^32^ that accounts for the spatial distances between cells. This demonstrates how our computational pipeline can readily accommodate alternative tools for deriving interaction strengths, depending on the specific needs and goals of the user. Importantly, our pipeline is modular and can be adapted to incorporate alternative toolkits, methods, or metrics for quantifying interaction strengths, depending on the context and specific needs (see Section 3).

In humans, *TGFβ*_1_, *TGFβ*_2_, and *TGFβ*_3_ are the three main isoforms of the transforming growth factor-beta (*TGFβ*) family. Each isoform initiates signaling by binding to specific surface receptors, leading to the formation of receptor-ligand complexes that in turn trigger intracellular cascades. *TGFβ*_1_, the most prominent and well-studied isoform in wound healing, first binds to the type II receptor (*TGFβR*2). This complex then recruits the type I receptor (*TGFβR*1) to form a heterotrimeric complex capable of initiating intracellular SMAD signaling:

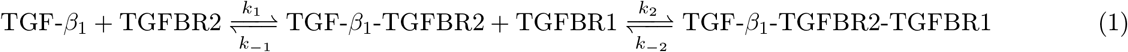

Likewise, *TGFβ*_3_ follows a similar receptor-binding mechanism:

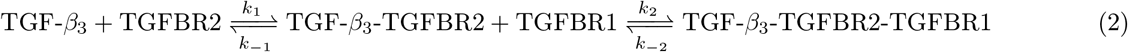

In contrast, *TGFβ*_2_ has low affinity for *TGFβR*2 and typically requires the presence of the co-receptor *TGFβR*3 (also known as betaglycan), which facilitates ligand presentation to *TGFβR*2 before the aformentioned binding cascade proceeds:

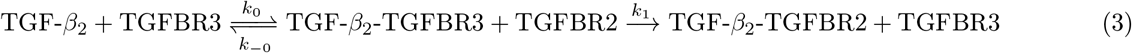

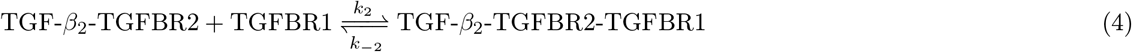

At steady state, such as during the wound remodelling phase, the net flux through each step of these binding cascades can be assumed to be equal (*i*.*e*, the formation and consumption rates of each intermediate complex are balanced).^33^ Therefore, to simplify the modelling of ligand-receptor dynamics within our reaction-diffusion framework, we focus only on the first binding event for each isoform, with corresponding binding rates ([*Ms*]^*−*1^)for isoforms 1,2, and 3 denoted as *k*_1_, *k*_2_, and *k*_3_, respectively. This allows us to represent ligand uptake via a single effective reaction step, while retaining the flexibility to extend to multistep kinetics in future refinements (see Section 3):

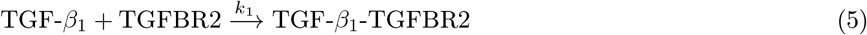

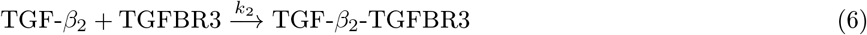

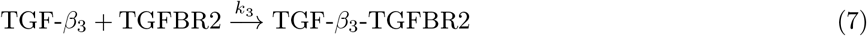

Considering the ligand-receptor interactions as defined in Equations 5-7, the resulting interaction strengths across the cell-type pairs and *TGFβ* isoforms as inferred from the trancriptomics data are shown in Figure 4.

**Figure 4.**
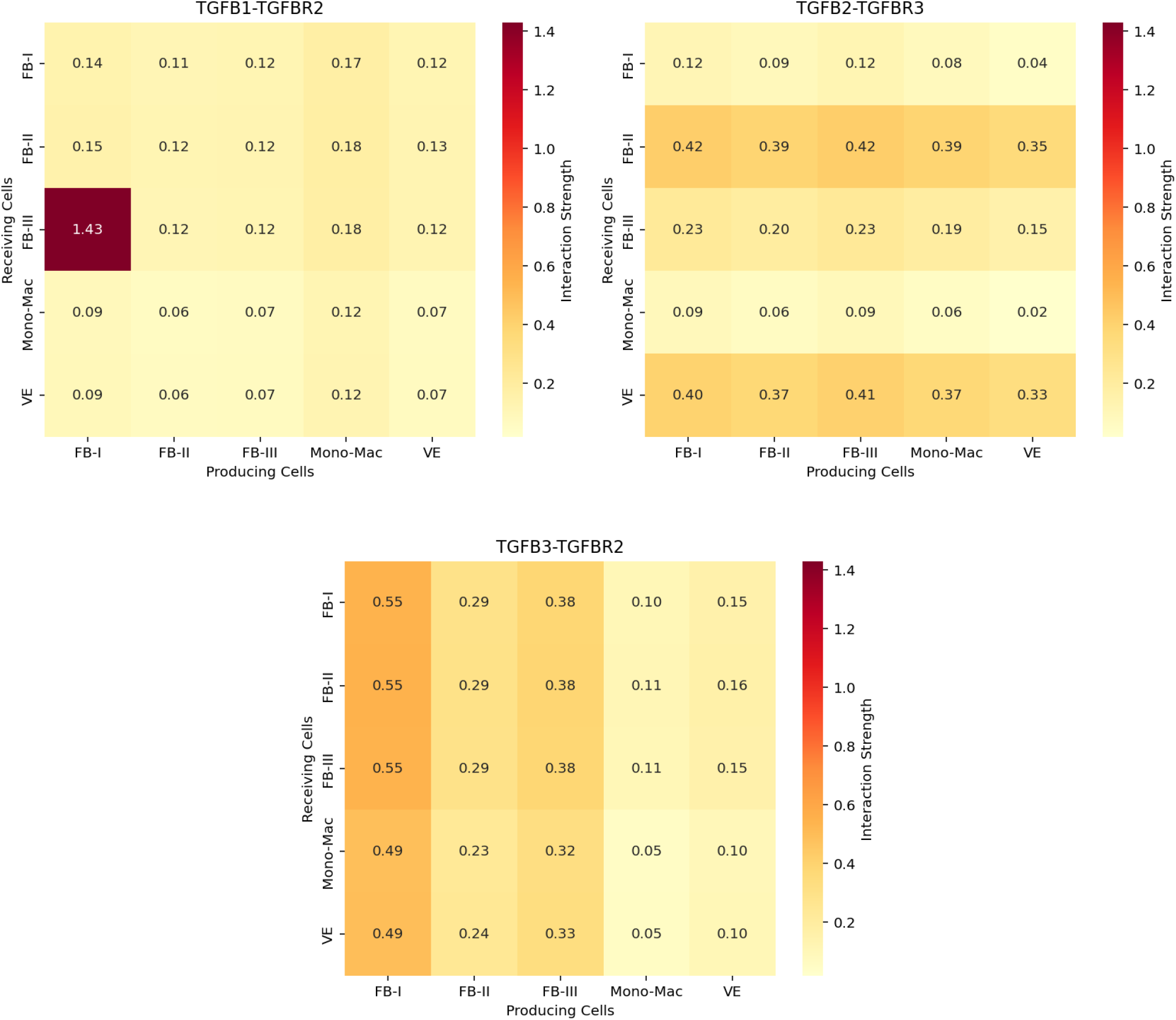
Interaction strengths between the cell types of interest across the three *TGFβ* isoforms, as inferred from CellPhoneDB analysis of the annotated scRNA-seq transcriptomic data for a normally healing tissue sample (donor 3, day 30 post-incision).^18^

### 2.2 Mathematical Model

The spatial distribution of a ligand (for example, a growth factor *g*) can be modelled via the following reaction-diffusion equation:

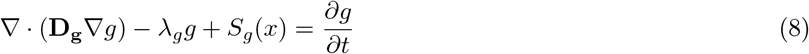

where, in the steady-state case, 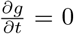. Here, **D**_**g**_ (in *m*^2^*s*^*−*1^) denotes the diffusivity tensor of the growth factor *g*; when assumed isotropic and spatially uniform, this simplifies to a scalar diffusion coefficient. The term *λ*_*g*_ (in *s*^*−*1^) represents the background decay rate of *g* in the domain, and *S*_*g*_(*x*) accounts for spatially-varying sources or sinks corresponding to production or absorption of the ligand. The source term *S*_*g*_(*x*) depends on the production by the different cell types *i* ∈ 𝒞 where 𝒞 denotes the set of all cell types, and on the local uptake of *g* due to binding to cell-surface receptors.

The choice between a continuum or particle-based formulation of Equation 8 in the computational pipeline depends partly on the nature and resolution of the available data. Methods like standard Visium spatial transcriptomics, which map gene expression to spatially localized spots encompassing multiple cells, and which can be combined with single-cell RNA sequencing to infer cell densities as carried out in Section 2.1.2, would be better suited for a continuous model. In this section, we adopt a continuum-based formulation, in line with the nature of the data (*i*.*e* the cell density distributions) extracted in Section 2.1.2. In contrast, when spatially resolved single-cell data is available, for instance, from technologies such as Visium HD, Multiplex Immunofluorescence (MIF), or Fluorescence In Situ Hybridization (FISH), particle-base formulations would be more suitable. In Appendix B, we derive the particle-based formulation and outline how it can be incorporated into the pipeline when the precise ceullar positions are available. Ultimately, the goal is to accommodate various spatial resolution data types while maintaining compatibility between the particle-based and continuum approaches.

In this study, we employ the finite volume (FV) method, which is well-suited for solving the type of conservation laws in Equation 8. Of course, other numerical methods may also be utilized, provided they follow the same core steps; that is, they proceed from discretizing both the equation and spatial domain, leading to a set of algebraic equations defined over control volumes (CVs) or finite elements, which can then be assembled into a system to solve for the spatial distribution of *g*. For consistency in units and to accurately reflect the geometry of the experimental data, we formulate the finite volume (FV) equations in three dimensions. This is necessary because certain parameters—such as molecular binding rates—are defined in volumetric units (*e*.*g*., [*M, s*]^*−*1^ = 10^*−*3^ m^3^/(mol · s)), requiring a 3D spatial context for proper dimensional alignment. Additionally, the tissue section under analysis, though visualized and discretized in 2D using a hexagonal grid, possesses a finite thickness in the third dimension. To account for this, we model each control volume (CV) as a hexagonal prism, enabling us to incorporate the sample’s depth while preserving the natural hexagonal tiling used in spatial transcriptomics platforms. Once the 3D formulation is established, we reduce the system to an effective 2D representation by collapsing the thickness dimension.

Using the divergence theorem on Equation 8, and assuming that the diffusivity tensor is isotropic and uniform, the divergence of the field (*D*_*g*_ ∇*g*) inside the CV can be related to the flux of the vector field through the CV closed surface boundary:

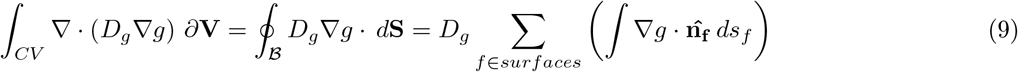

where 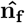 is the outward-directed unit normal vector to surface *f*. Given that Equation 8 is purely diffusion-based, we utilize a collocated arrangement for the FV method in which the values for *g* are stored in the FV centroids. Furthermore, since the PDE in Equation 8 is second-order in space, (at least) a second-order spatial discretization is warranted. Hence, the flux through the CVs can be formulated in terms of the value of ∇*g* at the edge centroids and can be written as:

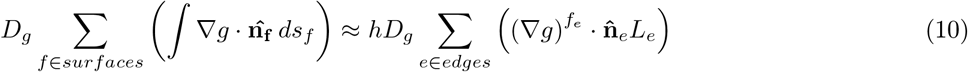

where *h* is the thickness of the CV prism, *L*_*e*_ is the length of the CV edge *e* in 2D, and 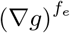 is the gradient of *g* evaluated at the centroid (*f*) of edge *e*. In a uniform orthogonal grid with no mesh non-orthogonality or skewness, the 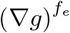 at the edge can be evaluated in terms of the value of *g* at the two centroids on the opposite sides of the edge, using the simple central difference scheme. This gives rise to 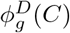 in Equation 11, which denotes the total diffusive flux of the growth factor out of CV *C*, in terms of the values of *g* at the centroid of the *C* centroid (*g*_*c*_) and those at the centroids of the neighbouring CVs (*g*_*i*_, *i* ∈ *N* (*C*)):

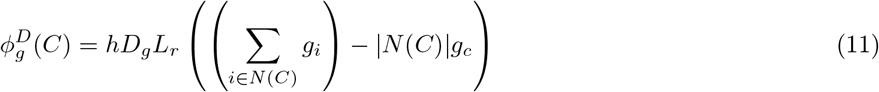

where *L*_*r*_ = *L*_*e*_*/L*_*C*_ and *L*_*C*_ is the distance between the centroids in the uniform mesh, and |*N* (*C*)| is the cardinal of *N* (*C*) (*i*.*e*., number of CVs adjacent to *C*). *S*_*g*_(*x*) in Equation 8 would represent any source or sink terms inside each CV due to ligand production or absorption by the different cell types, respectively. This can be written as:

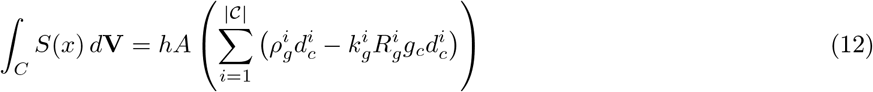

where *A* is the area of the control volume, 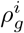 is the production rate (*mol/*(*s*.*cell*)) of ligand *g* by cells of type *i*, 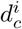 is the density of the cells of type *i* in the CV *C*, 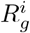 is the number of ligand *g*-binding receptors (*mol/cell*) on cell type *i* and 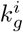 is the binding rate (*Ms*)^*−*1^ per cell between growth factor *g* and its corresponding receptor on cell type *i*. The definition is set up such that in the *lim*_*A*→0_, we recover the particle-based point source formulation in Appendix B. Note that in case there are multiple receptor types *j* binding ligand *g* on cell type *i*, the corresponding absorption rate inside the CV simply becomes 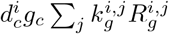.

Similarly, the decay rate within each CV can be formulated as:

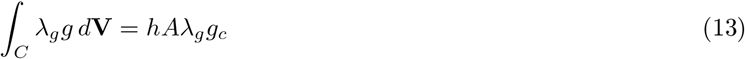

Equations 10, 12, and 13 can be divided by the thickness *h* and combined to form of a linear balance equation that accounts for the conservation of *g* within each CV *C* surrounded by neighbouring CVs *j* ∈ *N* (*C*) in the uniform orthogonal FV mesh:

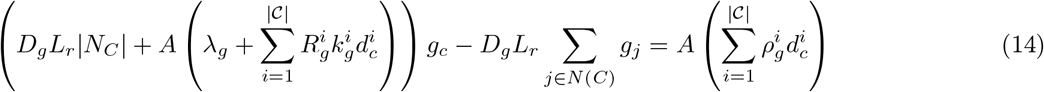

where the variables *A, L*_*r*_ and |*N*_*C*_| depend on the choice of the mesh and CV shape. Assuming no-flux Neumann boundary conditions (BCs), Equation 14 can be formulated for each CV and assembled to form a linear system of equations that can be solved for the growth factor concentrations across all CVs **g**:

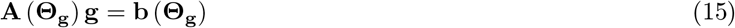

The entry values for matrix **A** and the column vector **b** in Equation 15 depend on the model parameter values, which are discussed in Section 3. The concentration of the growth factors at the receptor sites can then be used to calculate the ligand–receptor interaction strengths for the different ligand–receptor pairs on the various cell types across the tissue domain of interest. In the case of the continuous model, the interaction strength between cell type *l* secreting ligand *A* and cell type *r* expressing receptor *B* and can be approximated as:

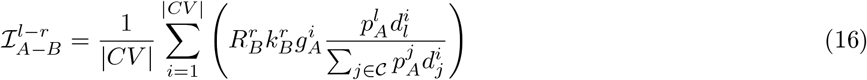

where |*CV*| is the number of CVs in the domain, 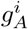 is the concentration of growth factor *A* in the centroid of 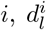 is the cell density of cell type *l* in CV *i*, and 𝒞 is the set of all cell types of interest. Equation 16 infers the contributions of a particular cell type in a particular spot to the total ligand binding in that spot. S

Since the relative scaling and offset between the model-derived and experimental interaction strengths has yet to be established, a correlation-based comparison, such as Pearson’s correlation coefficient *r*, can be employed to inform parameter calibration, since such measure is independent of absolute magnitudes; ultimately, the key objective is to identify the optimal model parameters that produce interaction strengths which, when compared pairwise, preserve the same relative ratios as those observed in the experimental data.

### 2.3 Parameter Initialization and Assembly

The first step in the parameter calibration framework involves constructing a prior distribution from which an ensemble of realizations can be initialized. To achieve this, we survey the current literature to either directly extract relevant parameter values or extrapolate them from available experimental and computational studies. Table 1 summarizes the key extracted parameters and outlines the rationale behind the assigned values and distribution.

**Table 1.** Initial parameter values for the *TGFβ* reaction-diffusion model. The values were collected from the literature and converted to SI units for dimensional consistency. This includes the cellular density of *TGFβ* receptors, which was converted from receptor count to number of moles by dividing by Avogadro’s constant. The notation ^A^ ← ^B^ indicates that the value(s) was extracted from the computational study in A based on the experimental results in B. *k*_1_, *k*_2_, and *k*_3_ correspond to the binding rates of *TGFβ* isoforms 1,2 and 3 to their corresponding *TGFβ* type 2,3 and 2 receptors, respectively. The LogUniform distribution of a variable *x* in the range [*a, b*] corresponds to uniform sampling in the *log*_10_ space such that to *log*_10_(*x*) ~ *U* [*log*(*a*), *log*(*b*)].

| Parameter<br>[Units] | Values<br>[Source(s)] | Initialization | Notes |
| --- | --- | --- | --- |
| $D$<br>$\left[ \times 10^{-11} \frac{m^2}{s} \right]$ | 1. 2.6 <sup>[34]</sup><br>2. 2.13 <sup>[35]</sup><br>3. 13 <sup>[36]</sup><br>4. 2.9 <sup>[10]</sup> | LogUniform<br>([2.13, 13.0]) | The same $D$ is assumed across all isoforms since they share similar molecular weights (25 kDa) and (80%) sequence identities.<br>The second value reflects the ligand’s interaction with the cartilage extracellular matrix (ECM), similar to what is observed in dermal tissues, leading to slower diffusion compared to free diffusion.<sup>36</sup> |
| $\lambda_g$<br>$\left[ \times 10^{-6} s^{-1} \right]$ | 1. 13.32 <sup>[37 ← 38]</sup><br>2. 4.10 <sup>[10 ← 21]</sup> | Uniform<br>([4.10, 13.3]) | The same $\lambda_g$ is assumed across all isoforms since they share similar molecular weights (25 kDa) and (80%) sequence identities.<br>The first value is based on a $TGF\beta$ life cycle of 48 hours used in the computational model<sup>37</sup> based on the experimental results<sup>38</sup>; we assume that this corresponds to a 90% decay of initial concentration in order to calculate the decay constant value.<sup>36</sup> |
| $k_1$<br>$\left[ \times 10^2 \frac{m^3}{s \cdot mol} \right]$ | 1. 5.1 – 5.7 <sup>[14, 15]</sup><br>2. 7.2 <sup>[13, 17]</sup><br>3. 11.6 <sup>[13 ← 16]</sup><br>4. 210 – 310 <sup>[15]</sup> | LogUniform<br>([5.1, 310]) | The larger range of values from the last source correspond to the experimental conditions in which the receptors were immobilized.<sup>15</sup> |
| $k_2$<br>$\left[ \times 10^2 \frac{m^3}{s \cdot mol} \right]$ | 15 <sup>[13 ← 39]</sup> | LogUniform<br>([5, 100]) | |
| $k_3$<br>$\left[ \times 10^2 \frac{m^3}{s \cdot mol} \right]$ | 1. 7 – 7.8 <sup>[17]</sup><br>2. 6.9 – 7.9 <sup>[40]</sup><br>3. 7.4 <sup>[13 ← 17]</sup><br>4. 18 <sup>[13 ← 16]</sup><br>5. 730 – 830 <sup>[15]</sup> | LogUniform<br>([6.9, 830]) | The larger range of values from the last source correspond to the experimental conditions in which the receptors were immobilized.<sup>15</sup> |
| $\rho_{fib}$<br>$\left[ \times 10^{-22} \frac{mol}{s \cdot cell} \right]$ | 1. 1.556 <sup>[36]</sup><br>2. 3.472 <sup>[41]</sup> | Uniform<br>([1.556, 3.472]) | The first value is from liver cells in a microfluidic chip, and converted to mols assuming a $TGF\beta$ molecular weight of 25kDa.<br>The second value is based on the reported production rates in scleroderma and normal dermal fibroblast cells. |
| $\rho_{mac}$<br>[$\times 10^{-22} \frac{mol}{s \cdot cell}$] | N/A | $0.5 \times Uniform$<br>([1.556, 3.472]) | Assumed to have a uniform distribution over half the range of $\rho_{fib}$, scaled by 0.5, since TGFβ secretion by fibroblasts is generally greater than that by macrophages during the remodelling phase of wound healing.<sup>42</sup> |
| $\rho_{EC}$<br>[$\times 10^{-22} \frac{mol}{s \cdot cell}$] | N/A | $0.2 \times Uniform$<br>([1.556, 3.472]) | Assumed to have uniform distribution over one fifth the range of $\rho_{fib}$, scaled by 0.2, since TGFβ secretion by ECs is generally the least compared to that by fibroblasts and macrophages during the remodelling phase of skin wound healing.<sup>42</sup> |
| $R_{fib}$<br>[$\times 10^{-20} \frac{mol}{cell}$] | 1.387-2.467 [41] | $Uniform$<br>([1.387, 2.467]) | Based on the reported number of receptors in scleroderma and normal dermal fibroblasts. |
| $R_{mac}$<br>[$\times 10^{-20} \frac{mol}{cell}$] | 0.0631-0.0640 [43, 44]<br>(for monocytes) | $LogUniform$<br>([0.0631, 1.329]) | The reported values correspond to monocytes; upon activation to macrophages, receptor numbers can rise significantly. This justifies the chosen starting distribution, which spans the lower monocyte values up to levels typical of fibroblasts and endothelial cells. |
| $R_{EC}$<br>[$\times 10^{-20} \frac{mol}{cell}$] | 0.997-2.193 [45] | $Uniform$<br>([0.997, 2.139]) | The range of values covers the number of TGFβ receptors under physiological and receptor expression-inhibited conditions. |

The LogUniform distribution used for some parameters in Table 1 yields a right-skewed profile that inherently favors smaller values. This choice is appropriate when literature-reported values tend to cluster at the lower end of the range, while still accommodating the full span of observed magnitudes with greater weighting on the more frequently reported smaller values.

Quantitative secretion rates of *TGFβ* per cell are not directly reported for macrophages or endothelial cells. However, M2 macrophages are well recognized as significant *TGFβ* producers during wound healing, whereas endothelial cells (ECs) contribute to a lesser extent, particularly in the remodeling phase. Accordingly, as an initial estimate, the production rate distributions for macrophages and ECs were scaled to 50% and 20% of the fibroblast baseline distribution in Table 1, consistent with the functional hierarchy *ρ*_*fib*_ *≥ ρ*_*mac*_ *≥ ρ*_*EC*_ described by Deng et al.^42^

In our analysis, each *TGFβ* isoform is treated as an independent ligand whose spatial distribution is governed by the reaction-diffusion equation described in Section 2.2, giving rise to a concentration field for each isoform. The concentration fields are combined with selected model parameters to compute the interaction strengths across the cell types for each isoform, according to Equation 16. The interaction strengths for each isoform can then be expressed in the form of an *n × n* matrix (where *n* is the number of the cell types of interest); this format mirrors the structure of experimentally inferred interaction strengths derived from high-throughput genetic sequencing data, as illustrated in Figure 4.

There are currently no pharmacological tools available to easily distinguish between *TGFβ* receptor subtypes, either via ligand-binding assays or through receptor-specific agonists or antagonists.^19^ Consequently, the distribution of receptor subtypes cannot be readily determined. Likewise, the reported *TGFβ* production rates do not differentiate among the three isoforms of the ligand. To account for this, our model randomly partitions the total reported *TGFβ* production rates and receptor abundances for each cell type among the three isoforms, introducing additional degrees of freedom to the parameter space. The parameters are then grouped into three distinct sets, one per isoform. For each realization, the diffusion coefficient (*D*) and degradation rate (*λ*_*g*_) are assumed to be shared across all isoforms, reflecting the high similarity in molecular profiles among the *TGFβ* isoforms. This assumption helps avoid unnecessary expansion of the parameter space, thereby reducing the risk of overfitting. Note that, in the specific case of analyzing data from normal skin wound healing at day 30 post-incision,^18^ three distinct fibroblast clusters were identified. Each cluster was treated as a separate cell type, with the corresponding parameter values sampled from the fibroblast-specific distributions listed in Table 1.

Using each isoform-specific parameter set, the growth factor concentration field is computed by solving the finite volume (FV) linear system described in Equation 15. The resulting spatial distributions are then used to calculate the corresponding interaction strength matrices between cell types according to Equation 16. Finally, the interaction strengths across all isoforms and cell types are concatenated, flattened, and compared against bioinformatics-derived interaction strength data. The Pearson correlation coefficient is computed to quantify the agreement between model-predicted and data-inferred interaction patterns. The initialization, partitioning, and reassembly steps used in this parameter assembly framework are illustrated in Figure 5.

**Figure 5.**
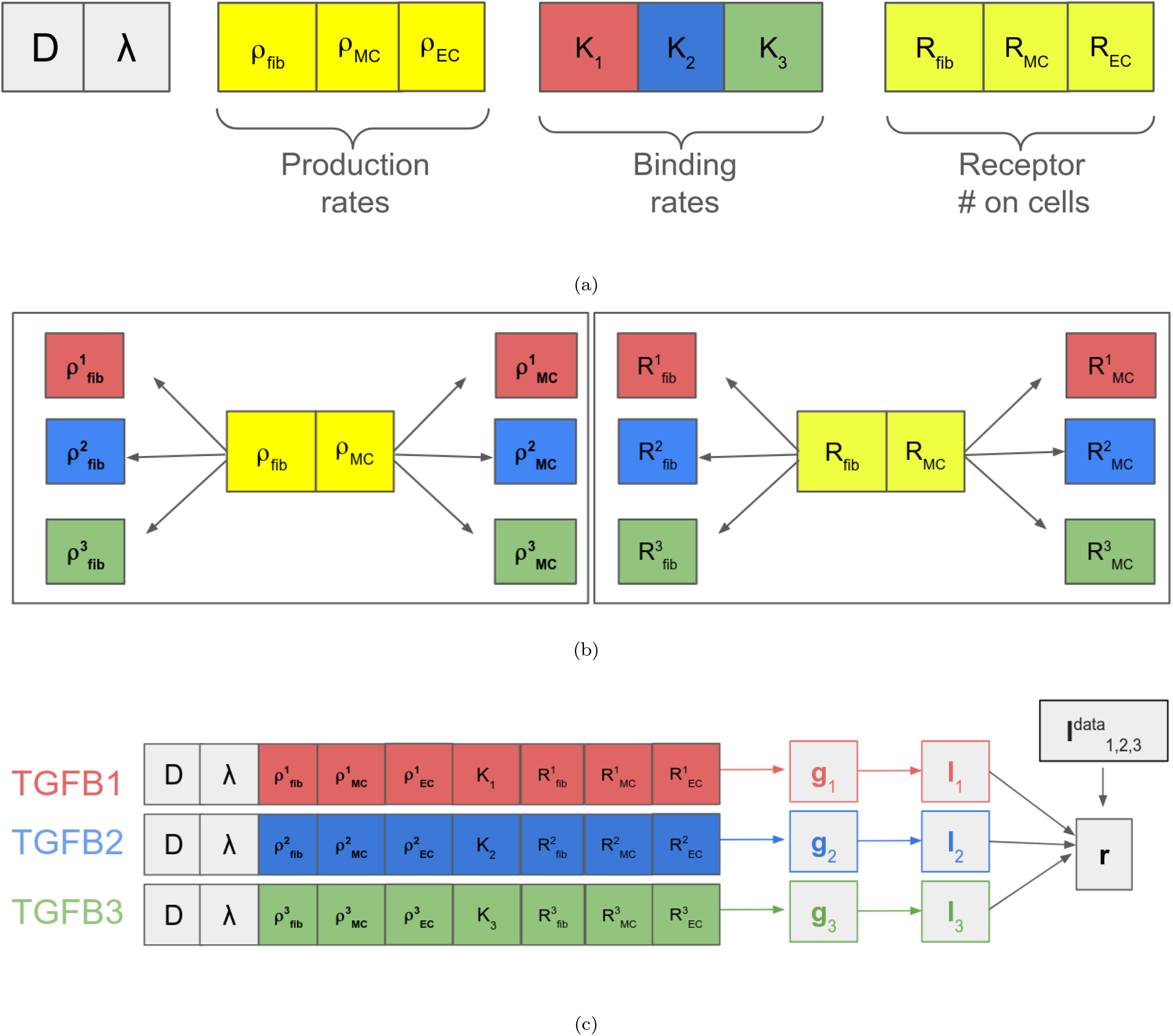
The parameter assembly framework for a particular parameter realization when modelling the reaction-diffusion dynamics of the *TGFβ* isoforms across the five cell types (the three fibroblast subtypes are represented here by one fibroblast type for brevity). A multidimensional parameter realization is sampled from the prior distribution (a). The production rates and number of receptors per cell are randomly partitioned across the three isoforms; the figure only shows the partitioning for the fibroblasts and macrophages, with similar portioning conducted for the EC cell type (b). The parameters are reassembled such that each isoform gets its own set of parameters. The *D* and *λ*_*g*_ parameters are assumed to be common across all isoforms. For each isoform, the FV linear system is solved to obtain the ligand concentration field, from which the model interaction strength matrix is obtained. The interaction strengths across the cell types and isoforms are then compared to the bioinformatics-derived interaction strengths data to obtain the Pearson’s correlation coefficient as a measure of agreement between the two datasets.

As mentioned in Section 2.2, because the relative scaling and offset between the model-derived and experimental interaction strengths are unknown (and likely depend on the specific bioinformatics tool used for inference), we employ a correlation-based approach to quantify agreement between the two. Specifically, both the model and data interaction matrices are flattened into vectors and compared using Pearson’s correlation coefficient (*σ*). This metric is invariant to scale and offset and captures linear relationships between the two sets of interaction strengths. While Pearson’s correlation is used here, alternative monotonic (but not necessarily linear) relationships may also be considered. Future work can explore the precise relationship between model-derived interaction strengths and those inferred by specific bioinformatics pipelines (see Section 3).

### 2.4 Parameter Calibration

In this section, we outline the development of a parameter calibration framework that integrates the bioinformatics-derived data on the spatial distribution of cell densities and interaction strengths (see Section 2.1) with the reaction-diffusion numerical modelling approach described in Section 2.2. The first step involves reconstructing the spatial in silico discretization mesh used in the finite volume (FV) numerical scheme, based on the spatial positions of the Visium spots arranged on a hexagonal grid. Given that the resulting computational mesh is uniform, orthogonal, and non-skewed, it can be fully characterized using two key pieces of information extracted from the Visium spatial transcriptomics layout.

First, the geometric properties of the mesh must be extracted; in particular, the control volume (CV) area *A* and the ratio of the edge length to the centre-to-centre distance *L*_*r*_ used in Equation 14 can be obtained from the knowledge of hexagon CV diameter (84.5*µm*). Second, the connectivity between adjacent CVs must be determined. This can be captured by a square boolean adjacency matrix where each *i, j* entry indicates whether CVs *C*_*i*_ and *C*_*j*_ are directly adjacent, and where adjacency is defined via the indicator function ℐ(||*C*_*i*_ − *C*_*j*_ || ≤ *L*_*c*_ + *ϵ*), with *L*_*c*_ representing the centre-to-centre distance between neighbouring hexagonal CVs and *ϵ* a small numerical tolerance.

These mesh-related components, along with the cell density maps for each cell type and the experimentally inferred interaction strength matrices, are treated as fixed inputs throughout the parameter calibration process. In contrast, the model parameters listed in Table 1 and shown in Figure 5 are treated as variable parameters subject to calibration. Below, we discuss several relevant parameter calibration schemes, highlighting the advantages and limitations of each, and describing how they can be sequentially combined within a hierarchical calibration framework that leverages the strengths of each approach while mitigating their respective weaknesses.

#### 2.4.1 Gradient Descent

The core principle behind using gradient descent is that the inputs (*i*.*e*., parameter realizations) are passed through a sequence of well-defined, differentiable mathematical operations to compute the final metric; in this case, it is the Pearson correlation coefficient (*r*). This structure enables the use of back propagation to trace how changes in the inputs affect the output, allowing us to iteratively update the parameters and minimize the discrepancy between the model-predicted and data-inferred interaction strengths.

To implement this, we first define a fitting loss function to be minimized. A natural choice in this context is:

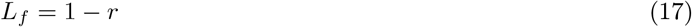

To prevent overfitting and to constrain the parameters within biologically plausible ranges, we incorporate a regularization term in the loss function. For each parameter *i* ∈[1, *d*] in the standardized parameter space, where *d* is the total number of tunable parameters, we define an acceptable range *R*_*i*_ = [*a*_*i*_, *b*_*i*_]. These bounds are informed by, or slightly broader than, the prior parameter ranges reported in Table 1. We then define a regularization function *L*_*R*_ that penalizes parameters falling outside their respective allowed intervals:

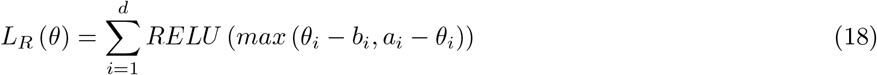

The total loss function is given by a weighted sum of the fitting loss and the regularization loss, with a hyperparameter *λ* controlling the trade-off between fitting accuracy and regularization:

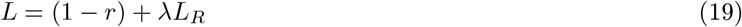

Note that this *λ* is distinct from the decay rate of the ligand *λ*_*g*_. Figure 6 demonstrates how the input parameters (a simplified representation using only two cell types) are propagated through a sequence of mathematical operators to obtain the final loss function.

**Figure 6.**
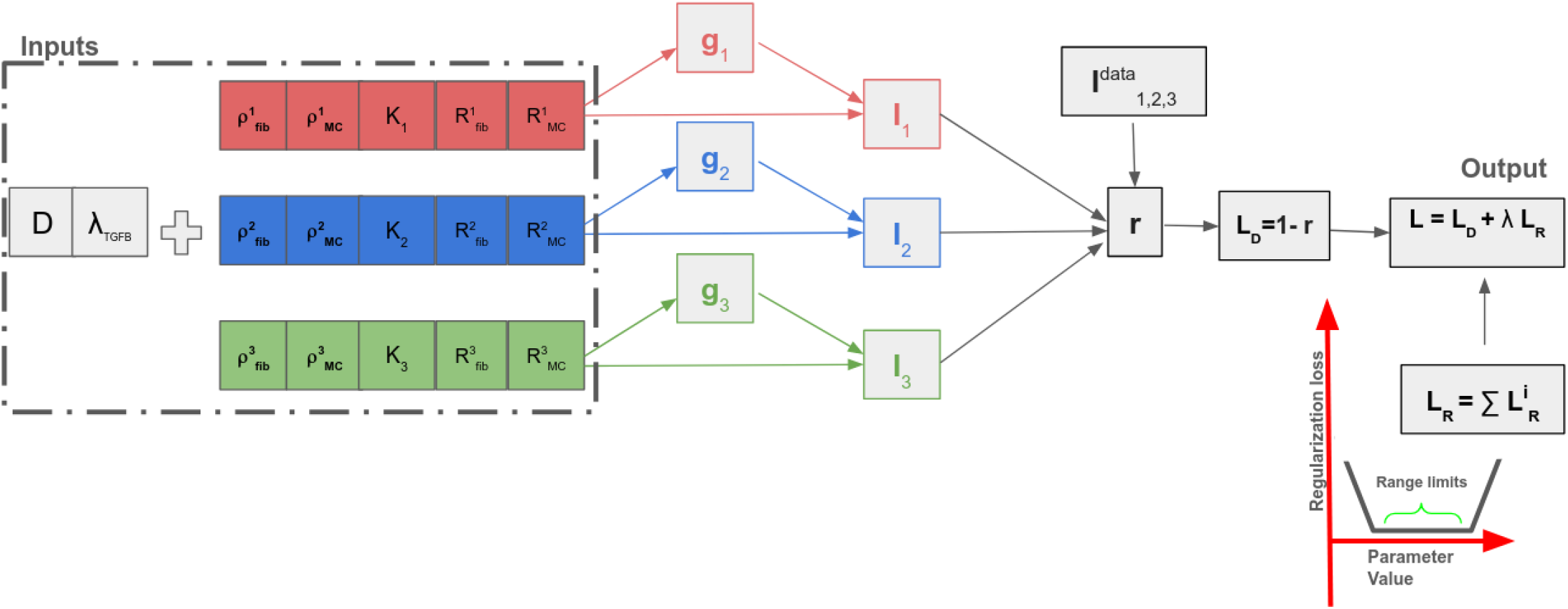
A simplified schematic demonstrating how the input parameter realizations are mapped through a series of mathematical differentiable operators to obtain the total loss function, which is comprised of the weighted sum of the fitting and regluarization loss functions. Using gradient descent algorithms, the effects of the parameters on the loss function can be traced back to update the parameters and minimize the loss function.

Next, applying the chain rule, we can map the dependence of the loss function *L* on any parameter Θ_*i*_ through the intermediate interaction strengths describing the cross-talk between the secreting cell type *l* and the receiving cell type *r*, across all *TGFβ* isoforms *j* = 1, 2, 3:

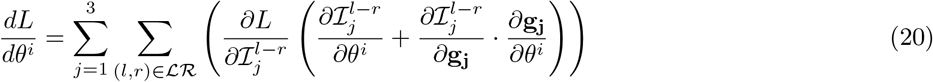

Note that in the case of the TGF*β* model, as only the parameters *D* and *λ*_*g*_ are shared across the isoforms, the partial derivatives of the loss with respect to the remaining parameters in Equation 20 are relevant to one isoform and do not require summing over all isoforms. Also note that the growth factor concentration field for each *j* ligand (isoform) **g**_*j*_ is obtained from solving the linear system of Equations in 15. In this system, both the LHS matrix **A**_*j*_ (**Θ**) and RHS vector **b**_*j*_ (**Θ**) depend on the parameters **Θ**, implying that **g**_*j*_ carries an implicit dependence on the parameters as well. To compute the Jacobian of **g**_*j*_ with respect to the parameters **Θ**, we can implicitly differentiate Equation 15:

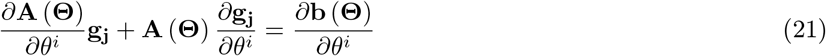

Rearranging the terms, we obtain a linear system to solve for the parameter derivative of the concentration field 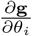:

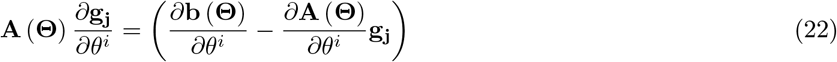

Note that both the original system of Equations in 15 and the Jacobian system in 22 involve the construction of the same matrix **A** (Θ), which can be leveraged for computational efficiency to solve for both systems of equations at every iteration of the gradient descent algorithm by reusing the LU factorization of **A** (**Θ**). Implementation-wise, the Jacobian matrix *∂***g**_*j*_*/∂***Θ** can be computed via a custom autograd function in Python, and then incorporated into back propagation chain of Equation 20 which in turn can be implemented within a gradient descent framework like PyTorch.^46^

Next, we outline the key advantages and limitations of applying a gradient descent algorithm, particularly in the context of parameter calibration for reaction-diffusion models. As previously discussed, one major advantage of this approach lies in the fact that the entire computational pipeline, from input parameters to the final loss metric, is explicitly defined through a sequence of differentiable operations. This enables the precise backpropagation of gradients, allowing the effects of changes in the input parameters to be directly mapped onto the loss function, and thereby facilitating targeted fine-tuning of the input parameters. In addition, the inclusion of a regularization term helps constrain the parameter values within biologically plausible ranges, reducing the risk of overfitting.

However, there are several challenges to consider. In a moderately high-dimensional parameter space such the TGF*β* model, different parameters exhibit varying degrees of sensitivity or constraint: some parameters must lie within narrow bounds to yield a low fitting loss, while others permit broader variation. This heterogeneity suggests that using a single, uniform learning rate across all parameters may be suboptimal. Instead, the relative learning rates across the parameters should ideally be adapted to reflect their respective relative degrees of constraint. Moreover, the optimization landscape in such models is likely to be non-convex and may contain multiple local minima. As a result, convergence to a global minimum is not guaranteed. This issue can be partially mitigated by initializing multiple optimization runs, or ‘seeds’, at different starting points that span the parameter space. However, each gradient descent run can be computationally expensive, and increasing the number of seeds further compounds the computational cost.

Considering these factors, an effective strategy would be to carefully select a small number of well-informed initialization seeds that are likely to be near regions of low loss and that collectively provide good coverage of the parameter space. The optimization performance can also be improved by assigning parameter-specific relative learning rates that reflect the relative degrees of constraint or sensitivity among the parameters.

#### 2.4.2 The Rejection Sampler Algorithm

Next, we look at Approximate Bayesian Computation (ABC) techniques to calibrate the ligand reaction-diffusion model parameters. ABC is a class of computational methods rooted in Bayesian statistics that can be used to estimate the posterior distributions of model parameters without the explicit evaluation of the likelihood function, which is often computationally intractable and/or too costly to evaluate.^47^ Let **Θ** be a parameter vector to be estimated. Given some prior distribution *π*(**Θ**), the goal is to approximate the posterior distribution that integrates knowledge from the available experimental data: *π*(**Θ**|*x*) ∝ *f* (*x*|**Θ**)*π*(**Θ**), where *f* (*x*|**Θ**) is the likelihood of obtaining the data *x* given the parameter space **Θ**.^47^ In ABC, the likelihood is replaced by a comparison between simulated and experimental data, using Monte Carlo simulations to explore parameter space. In our application, the comparison is based on model-predicted interaction strengths derived from the simulated ligand concentration fields. The final posterior distribution thus reflects the combined information from model dynamics, prior knowledge, and experimental measurements.

The simplest ABC algorithm would be the ABC rejection sampler,^48^ the general steps for which in the context of calibrating the reaction-diffusion model are summarised below:

1. Sample candidate parameter vector **Θ**\* from some proposal (prior) distribution *π*(**Θ**). In our case, the prior distribution can be constructed by screening the experimental and computational data from the relevant literature as described in Section 2.3.
2. Use the parameter vector **Θ**\* to simulate a dataset *x*\* from the dynamic model, In our case, this would correspond to the concentration fields of the ligand(s). The dataset *x*\* can be used to compute some secondary output *y*\*, such as the interaction strengths.
3. Compare the simulated output *y*\* with the corresponding experimental data (*x*_0_ or *y*_0_), using a scoring function or similarity measure and threshold *ϵ*. We use the Pearson’s correlation coefficient *r* as our similarity measure and accept **Θ**\* if *r*(*y*_0_, *y*\*) *≥ ϵ*, otherwise reject.
4. Return to step (1) and repeat for a predetermined number of trials or until the desired number of samples is achieved.

The model outputs are thus partitioned into two sets of accepted and discarded realizations, and the accepted set is mapped back into the input parameters^49^ to provide an empirical approximation to the posterior distribution *f* (**Θ**|*x*).

The ABC rejection sampler offers several advantages. It is straightforward to implement computationally and, because steps 1–3 of the algorithm are independent, it is embarrassingly parallelizable,^47^ for instance, across multiple cores within a high-performance computing (HPC) cluster. Moreover, by comparing the prior and posterior parameter distributions, one can gain valuable insight, at no additional computational cost, into parameter identifiability, the degree of constraint imposed by the data, and the sensitivity of model outputs to parameter variations; see^47^ for an example.

Unlike gradient descent, the ABC rejection sampler does not provide a mechanism to map discrepancies between model outputs and data-derived outputs back onto the inputs for precise parameter fine-tuning. Furthermore, the acceptance rate of the ABC rejection sampler can be very low when prior distributions are poorly informed or substantially different from the posterior distributions.^47, 50, 51^ In our model, while some priors can be constructed on physiological grounds from experimental data (see Section 2.3), reliable prior approximations are not available for other parameters (see discussion in Section 1). This issue is exacerbated in multidimensional parameter spaces, where inefficiencies can arise due to mismatches between initial sampling and the target posterior distribution defined by the acceptance criterion *r*(*y*_0_, *y*\*) ≥ *ϵ*. For example, a parameter may individually exhibit a high likelihood, but if combined with low-likelihood values of other parameters, the low joint posterior probability will result in poor acceptance criterion of the former.^52^ Overall, when prior knowledge is limited for some parameters, the ABC rejection sampler can be viewed as a preliminary step that produces an improved prior distribution better aligned with the ex vivo transcriptomics data, while simultaneously providing useful insight into parameter constraints and identifiability.

#### 2.4.3 Markov Chain Monte Carlo ABC schemes

When the posterior distribution is far from the prior, sampling directly from the prior in the ABC rejection sampler is often inefficient due to low acceptance rates.^50^ To address this, Markov Chain Monte Carlo (MCMC) methods such as the Metropolis-Hastings algorithm can be employed. MCMC constructs a Markov chain whose stationary distribution corresponds to the target posterior *π*(**Θ**|*x*). For likelihood-free simulations, Marjoram et al.^50^ introduced an MCMC scheme for generating observations from a posterior distribution without the need for the likelihood calculation, which we refer to here as ABC MCMC. The ABC MCMC scheme proceeds as follows^50, 52^:

1. Initialize **Θ**_*i*_, *i* = 1.
2. Generate a candidate parameter realization **Θ**\*~*q*(**Θ**|**Θ**_*i*_) where *q* is the proposal density, or transition kernel. Common transition kernels include Gaussian distributions with a mean of **Θ**_*i*_ and some specified variance.
3. Generate the dataset *x*\* from the model and **Θ**\* and use it to compute the model output *y*\*.
4. Compare the simulated output, *y*\*, with the experimentally derived output *y*_0_ using a similarity measure (*r*) and an acceptance threshold *ϵ*.
5. If *r*(*y*_0_, *y*\*) *< ϵ*, go back to step 2, otherwise set and accept **Θ**_*i*+1_ = **Θ**\* with probability: 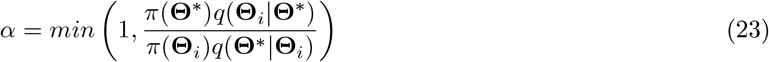

otherwise set **Θ**_*i*+1_ = **Θ**_*i*_.
6. if *i < N* where N is the prescribed Markov Chain length, increment *i* = *i* + 1 and go to step 2.

Thus, ABC-MCMC algorithms generate a sequence of serially and highly correlated samples^50^ which meet the criteria *r*(*y*_0_, *y*\*) ≥ *ϵ* after convergence is achieved. The major difference between the conventional and the ABC MCMC methods is that the former requires an explicit definition of the ratio of the likelihood functions to calculate the transition probability upon transition from **Θ** to **Θ**\*:

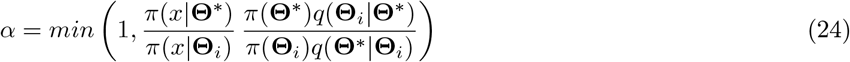

while in the latter case, if both **Θ** and **Θ**\* result in successful realizations (*r*(*y*_0_, *y*\*) ≥ *ϵ*), and, assuming that the perturbation in the parameter values is small, the likelihood ratio is approximated as 1.

When the prior distribution is considerably different than the posterior distribution, the ABC-MCMC algorithm can deliver substantial increases in acceptance rate over ABC rejection sampler algorithm, as demonstrated in the case studies by Majoram et al.^50^ One disadvantage of such MCMC algorithms, however, is that the resulting Markov chain samples are serially correlated, which, combined with potentially low acceptance probabilities or poorly chosen proposal distributions, can result in long chains that mix slowly or become trapped in low-probability regions.^47, 51, 52^

#### 2.4.4 Sequential Monte Carlo Filtering AMC schemes

In theory, generating a large number of uncorrelated and independent realizations (or particles) can help prevent Markov chains from becoming trapped in low-probability regions. Moreover, by starting from a broad prior distribution with a relatively loose error tolerance and gradually tightening the tolerance, ABC algorithms can more effectively explore high-dimensional parameter spaces without getting stuck in realizations of local extrema probabilities.^47^

To this end, Sequential Monte Carlo (SMC) ABC methods were introduced by Sisson et al.^51^ as an improvement over the simple ABC rejection sampler. In SMC-ABC, a sequence of particle populations is generated, each evolving through intermediate distributions that transition smoothly from the prior toward the target posterior. Examples of this progressive narrowing of posterior distributions in practice can be found in ^52, 53^.

At each iteration *t*, the acceptance threshold *ϵ*_*t*_ is smaller than in the previous iteration (*ϵ*_*t*_ *< ϵ*_*t−*1_), ensuring that the region of parameter space explored at iteration *t* is a subset of that explored at *t* − 1. Conceptually, this can be viewed as a sequence of rejection-sampling steps where the posterior from one iteration serves as the prior for the next. The gradual tightening of *ϵ*_*t*_ promotes efficient convergence to the posterior distribution.^47, 51, 52^ Starting from a prior distribution *π*(**Θ**), the general ABC-SMC algorithm proceeds as follows ^47^:

1. Initialize *ϵ*_1_, … *ϵ*_*T*_. Set the population indicator t=0.
2. set the particle indicator i=1.
3. Sample **Θ**\* from the previous population 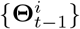 with weights *w*^*t−*1^ and perturb the particle (according to some perturbation Kernel) to obtain **Θ**\*\*~ *K*_*t*_ (**Θ**|**Θ**\*). If *π*(**Θ**∗∗) = 0 repeat the sampling. If If t=0 then sample **Θ**\*\* directly from the prior *π*(**Θ**).
4. Simulate a candidate dataset (*e*.*g*. the ligand concentration field) from the obtained parameter realizations *x*\* *~ f* (*x* | **Θ**\*\*), calculated the output *y*\* (*e*.*g*. the interaction strengths) and calculate the corresponding simmilarity statistic (*r*(*y, y*\*)); if *r*(*y, y*\*) *< ϵ*_*t*_, return to step 3.
5. Set 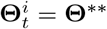 and calculate the weight for particle 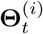:

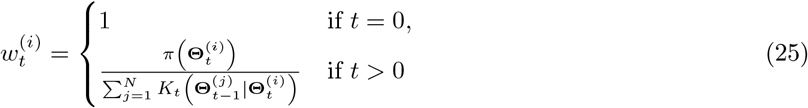

If i<N set i=i+1 and go to step 3.
6. Normalize the weights. If t<T set t=t+1 and go to step 2.

In this notation, particles sampled from the previous population before perturbation are marked with a single asterisk (*), while perturbed particles are marked with a double asterisk (**). When *T* = 1, this reduces to the ABC rejection sampler in Section 2.4.2.

In the original algorithm formulations,^47, 51^ both the thresholds *ϵ*_1_, …, *ϵ*_*T*_ and the perturbation kernel were specified a priori. In our implementation, these quantities are adaptively chosen: the threshold at iteration *t* is set to the 75th percentile of the accepted distances from iteration *t −* 1, i.e., *ϵ*_*t*_ = *q*_0.75_(*ϵ*_*t−*1_), and the perturbation kernel for the standardized variables is Gaussian: 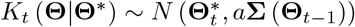 where **Σ** (**Θ**_*t−*1_) refers to the covariance matrix from the previous population and *a* is some scaling constant (set here to 1.2).

The gradual convergence towards the target distribution across sample populations in SMC schemes can yield a more efficient and targeted approach for reaching the optimal distribution than rejection sampler schemes. The generation of multiple independent particles at each population also enhances robustness against becoming trapped in low-probability regions compared to MCMC algorithms. Moreover, the SMC algorithm is generally less sensitive than the rejection sampler to poorly informed prior distributions (though not completely insensitive, as it still relies on the prior for initialization and particle weight allocation in Steps 3 and 5, respectively). In addition, similar to the rejection sampler algorithm in Section 2.4.2, the sampling and processing of multiple particle realizations within each population (Steps 3–5) can be parallelized. Furthermore, as we have demonstrated, the algorithm can be reformulated in an adaptive form, in which the thresholds *ϵ*_1_, …, *ϵ*_*T*_ and the perturbation kernel *K*_*t*_ (**Θ** | **Θ**\*) are tuned dynamically based on the progress and behaviour of the algorithm. Finally, as with the rejection sampler in Section 2.4.2, additional insight into the degree of constraint for each parameter can be obtained at no extra cost through retrospective analysis of the parameter distribution evolution across populations.^47^

On the other hand, SMC algorithms become less practical for calibrating parameters in high-dimensional spaces due to the increased computational cost.^47^ The reason is twofold: first, a sufficiently large number of particles must be sampled at each population iteration to adequately cover and characterize the multi-dimensional parameter space. Second, in the case of adaptive algorithms devised and used in this study, it is necessary to compute the covariance matrix and construct a multivariate perturbation kernel after each population. However, covariance matrices can become less reliable and more prone to numerical instability as the number of parameters increases. In addition, while less sensitive than the rejection sampler ABC schemes to the choice of the prior distribution, the algorithm still requires the allocation of a close and well-informed prior distribution. Finally, as in the rejection sampler algorithm described in Section 2.4.2, there is no direct fine-tuning of input realizations based on the output, as is the case for the Gradient Descent algorithm in Section 2.4.1.

To this end, and in the specific context of calibrating the growth factor reaction–diffusion model parameters described in Section 2.3, the most effective approach is to employ the adaptive SMC algorithm described above after the allocation of an appropriate multivariate prior distribution (with a defined probability density function as required in step 5). To improve algorithm robustness and reduce computational cost, the dimensionality of the parameter space can be also decreased by fixing certain parameters a priori. Furthermore, as is the case with the rejection sampler scheme, analyzing the variances of the parameter distributions at both the initial and final populations can provide valuable insight into the degree of constraint associated with each parameter.

#### 2.4.5 Parameter Calibration: The Computational Pipeline

In Sections 2.4.1–2.4.4, we reviewed several relevant calibration schemes and outlined the advantages and limitations of each in the context of parameter calibration for the ligand reaction-diffusion model. In this section, we combine the rejection sampler and the SMC ABC schemes with gradient descent in a sequential/hierarchical manner into a single computational pipeline, aiming to leverage the strengths of each approach while mitigating their weaknesses. The parameter calibration pipeline is illustrated in Figure 7 and described in detail below, using the specific study case of *TGFβ* ligand reaction-diffusion parameter calibration.

**Figure 7.**
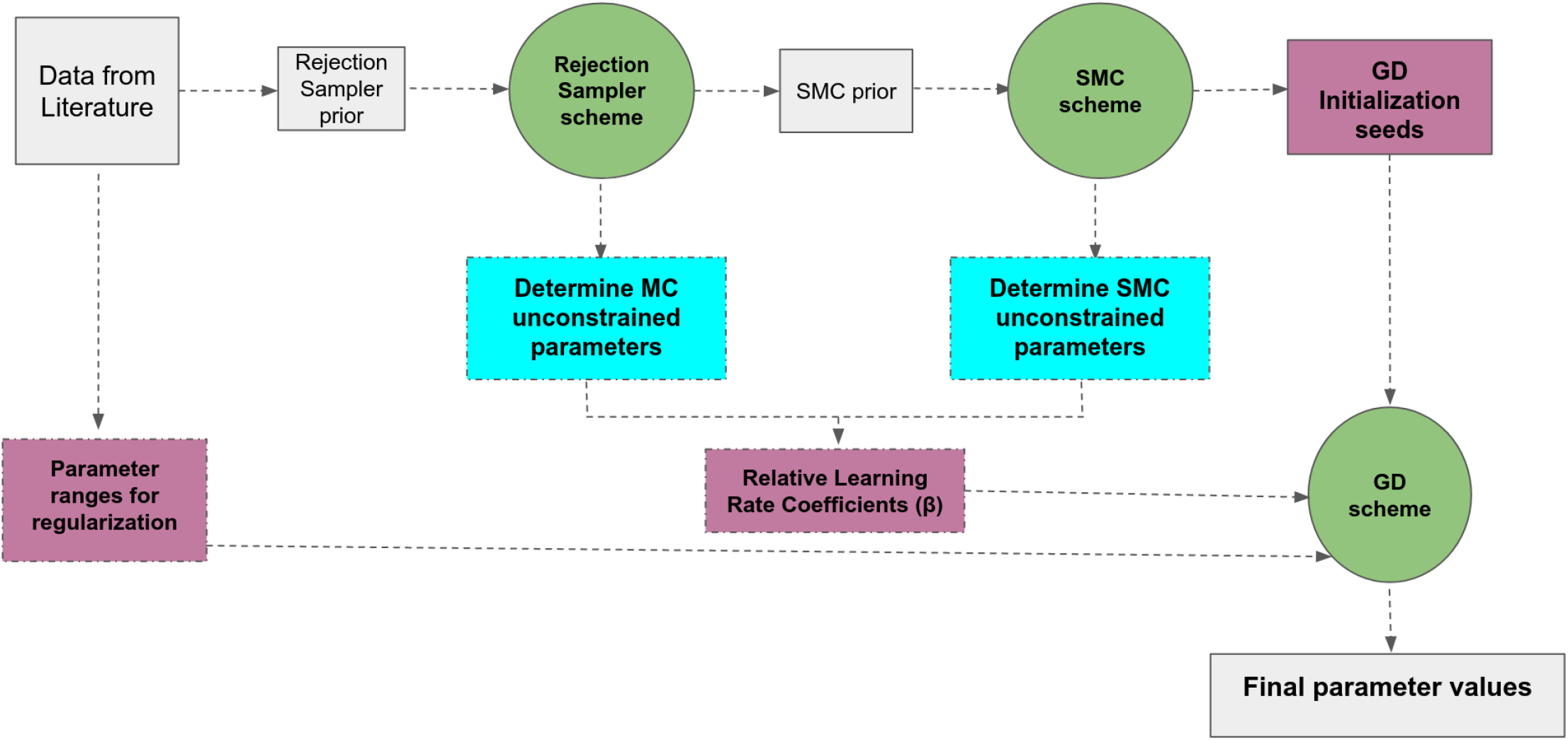
The computational pipeline for the parameter calibration of the ligand reaction-diffusion model from transcriptomics data.

First, the preliminary values for the parameters obtained from experimental and computational studies in the literature are used to construct a prior distribution for the rejection sampler algorithm described in Section 2.4.2. This constructed prior distribution is also used to extract the permissible ranges for parameter values to be employed in constructing the regularization loss function in Equation 18 for the gradient descent algorithm. The rejection sampler algorithm is then executed with *MC* = 1*e*6 independent parallelized runs and an acceptance threshold of *ϵ ≥* 0.2, yielding a reduced ensemble of accepted parameter realizations with *r* ∈ [0.2, 0.67], as shown in Figure 8.

**Figure 8.**
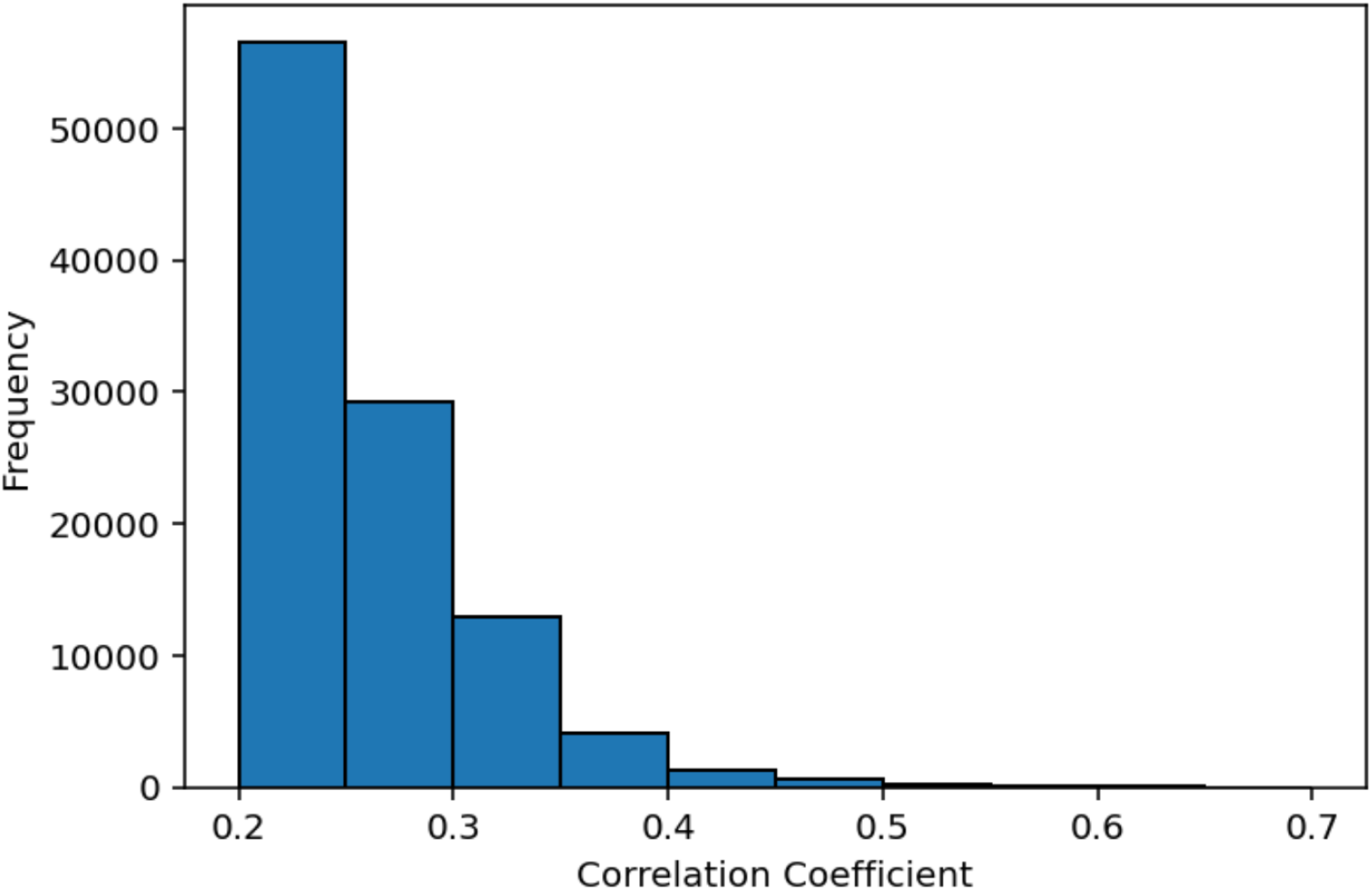
Histogram showing the distribution of the correlation coefficients for the accepted runs when applying the rejection sampler algorithm (*MC* = 1*e*6, *ϵ* = 0.2) to the specific study case of *TGFβ* ligand reaction-diffusion parameter calibration.

Furthermore, based on post-hoc sensitivity analysis of the parameters and their degree of constraint after the rejection sampler and SMC analyses, we can categorize the parameters into one of three categories based on their sensitivity and degree of constraint, which would in turn guide the allocation of the relative learning rates for the Gradient Descent algorithm, as described below.

For each parameter, the prior and posterior ensembles of realizations from the rejection sampler step are compared using the Kolmogorov–Smirnov (KS) test to determine whether the two continuous probability distributions are significantly different (*p <* 0.01). If the distributions are similar (*p ≥* 0.01), this indicates that the parameter does not significantly affect the acceptance rate and likely has an insignificant impact on the output, or that the prior is already close to the posterior and thus well-informed. Accordingly, these parameters are considered to have the least impact on the output and are assigned to the ‘Grade 1’ parameter category.

Alternatively, if the distributions are significantly different (*p <* 0.01), it indicates that the parameter has a noteworthy effect on the acceptance rate and model output, or that the prior distribution is far from the posterior. In total, 9 out of the 35 parameters from the TGF*β* model were assigned to the ‘Grade 1’ category. Figures 9(a) and 9(b) show examples of parameter distributions before and after applying the rejection sampler algorithm for parameters in ‘Grade 1’ category, where the distributions are statistically not different. Conversely, Figures 9(c) and 9(d) illustrate the corresponding changes for parameters whose prior and posterior distributions are statistically different. Notably, when simultaneously calibrating model parameters for the three TGF*β* isoforms, the parameters *D* and *λ*_*g*_ are always assigned to the ‘Grade 1’ category. This is likely because these two parameters are shared across all three isoforms and influence the ligand concentration fields and interaction strengths similarly across them.

**Figure 9.**
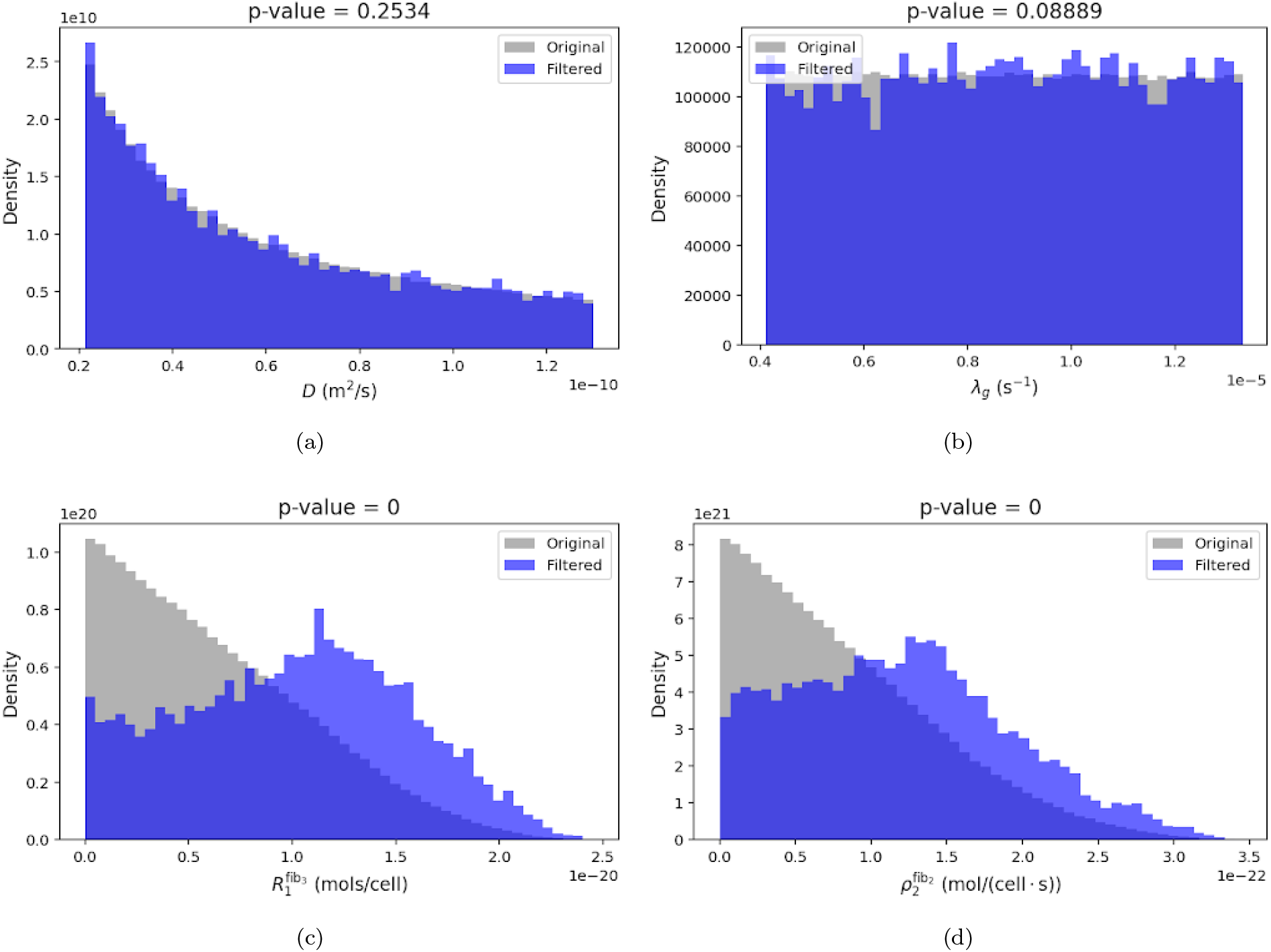
Examples comparing the prior (original) and posterior (filtered) distributions obtained from the rejection sampler scheme for the *TGF β* model. For parameters in the ‘Grade 1’ category, the prior and posterior distributions are not statistically different according to the KS test (top), whereas the two distributions for the remaining parameters are statistically different (bottom).

**Figure 10.**
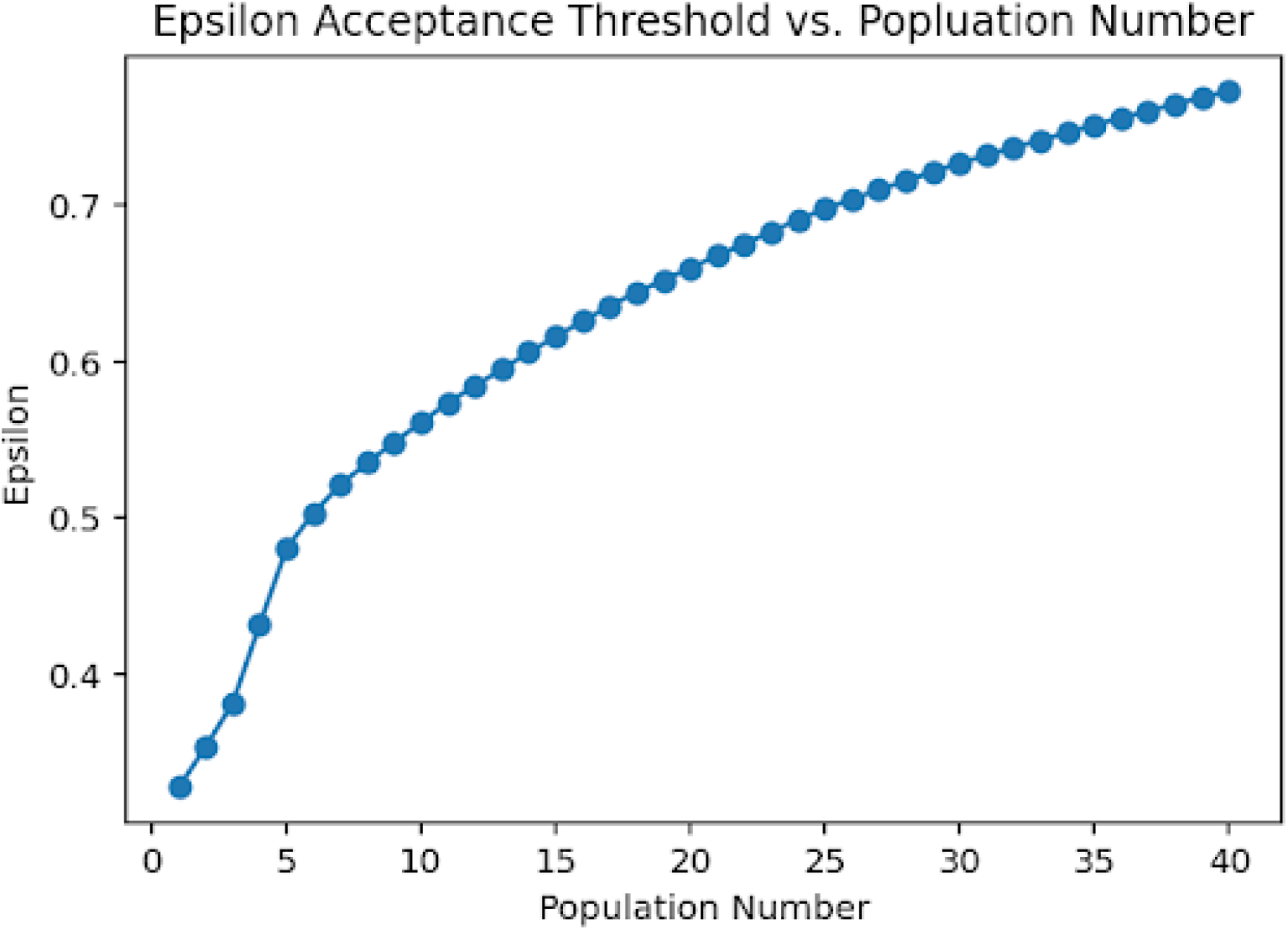
The evolution of the acceptance thresholds for the TGF*β* reaction-diffusion model throughout the SMC populations, where the acceptance threshold for each population is computed from *ϵ*_*t*_ = *q*_0.75_(*ϵ*_*t−*1_).

Next, the filtered ensemble of realizations obtained from the ABC rejection sampler algorithm is used as the prior distribution for the SMC algorithm, with the parameters designated to ‘Grade 1’ category from the preceding rejection sampler fixed (according to the maxmium likelihood estimator values) to reduce the parameter space dimensions. Since the SMC algorithm in Section 2.4.4 requires a prior distribution with an explicit probability density function (pdf), a posterior distribution, for example, a general multicomponent, multivariate Gaussian mixture model (GMM), is fitted to the filtered ensemble of realizations using <monospace>scikit-learn</monospace> in Python. In case of fitting a GMM, a Bayesian Information Criterion (BIC) analysis can be conducted to determine the optimal number of components. Alternatively, the BayesianGaussianMixture toolkit can be used, which starts with an upper bound on the number of components and shrinks away unimportant ones. Ultimately, the algorithm can accommodate multiple options for distribution fitting. Since the parameters span a wide range of scales and magnitudes, we standardize their values prior to distribution fitting, which takes place for the prior distribution (in order to obtain a pdf), as well as when computing covariance matrices for the adaptive perturbation kernel during each population iteration of the SMC algorithm. This standardization helps prevent scale dominance and numerical instability.

For the *TGFβ* model, the ABC SMC algorithm is executed with *N* = 2*e*4 particles and *T* = 40 populations. Figure 10 demonstrates the evolution of the acceptance thresholds for the TGF*β* reaction-diffusion model throughout the SMC populations, which reflects the gradual improvement in the correlation coefficients since the acceptance threshold for each population is computed from *ϵ*_*t*_ = *q*_0.75_(*ϵ*_*t−*1_).

As in the case of the ABC rejection sampler scheme, post-hoc analysis of the parameter distributions and their evolution can provide insight into output sensitivity to the parameters and their degrees of constraint at no additional computational cost. Figure 11(a) provides an example of a parameter for which the first and final population distributions are not significantly different; in such case these parameters are likely poorly inferrable. Conversely, Figure 11(b) provide an example for a parameter for which the distributions do significantly change between the first and last populations, accompanied by significant narrowing in the permissible range of parameter values, which means the parameters can likely be well-inferred.^47^

**Figure 11.**
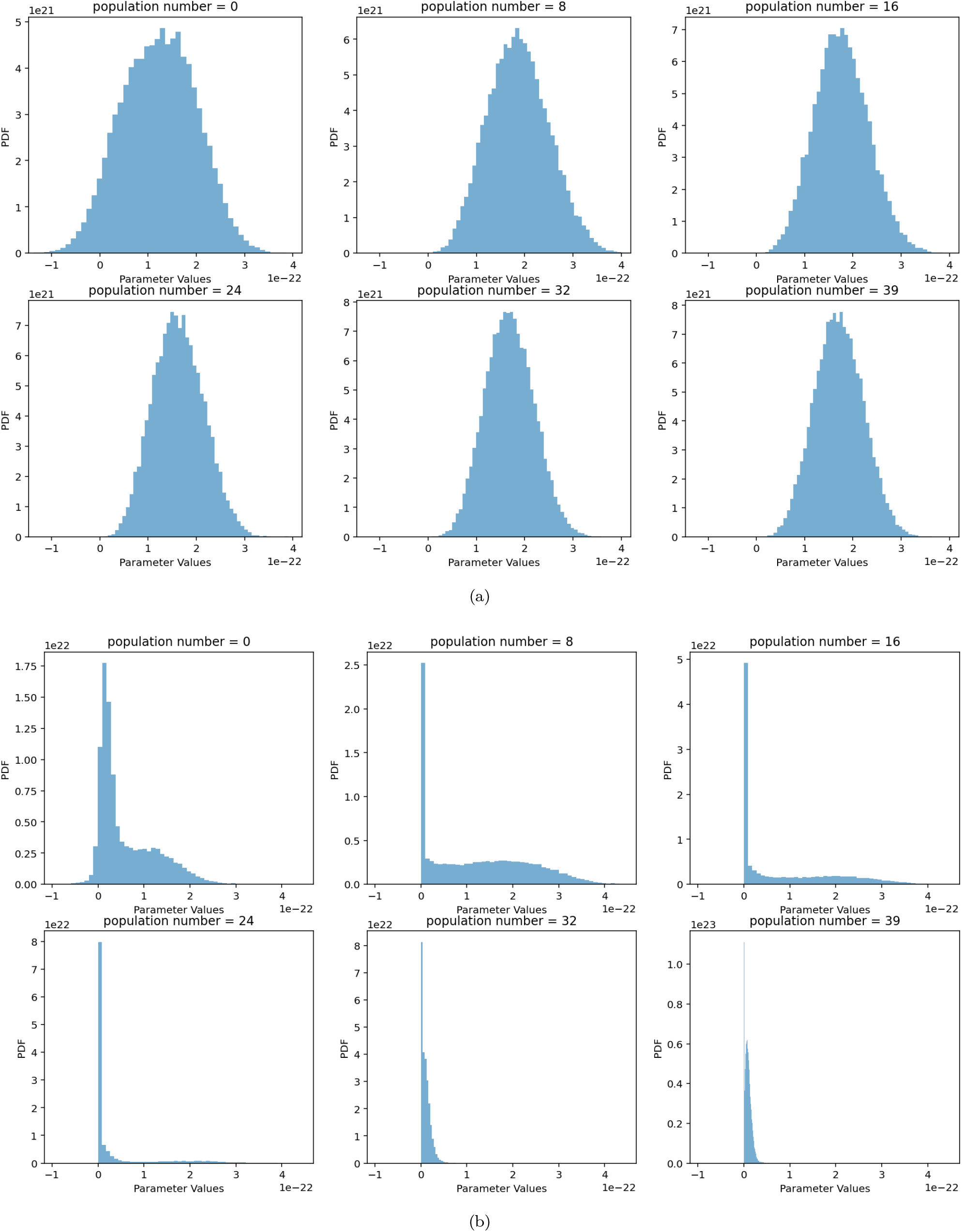
Examples tracing the evolution of the distributions across the SMC scheme for the TGF*β* model. For parameters in the ‘Grade 2’ category, the prior and posterior distributions are not different (top), whereas the final posterior distribution for ‘Grade 3’ parameters are significantly different and narrower than the starting prior distributions (bottom).

Note that, because the perturbation kernel models a Gaussian random walk, the perturbed parameter realization **Θ**\*\* may occasionally take on non-physiological negative values in the original (non-standardized) parameter space. Additionally, the algorithm may converge towards such non-physiological parameter values during the iterative evolution of the distributions if they improve the fit between the model and data-inferred interaction strengths. Therefore, after perturbation and prior to computing ligand concentration fields, the parameters are clipped to ensure they remain positive when converted to their original scale. This explains why, as some parameter distributions evolve across iterations, the accepted values may form a peak near zero in the original space, as evident in Figure 11.

In line with the method of sensitivity analysis by Toni et al.^47^ we can quantify the sensitivity of the system output to the parameters and their combinations by conducting a PCA analysis on the output of the SMC analysis, which is the ensemble of *N* particles from the last population. PCA provides an orthogonal transformation of the original parameter space into the new principal components (PCs), which are the eigenvectors of the corresponding covariance matrix Σ. Hence, each unit eignvector *PC*_*i*_ is a linear combination of the parameters in the original space, and the corresponding eigenvalue *λ*_*i*_ to each eigenvector *PC*_*i*_ (obtained via Singular Value Decomposition, SVD) is proportional to the amount of variance *var* (*PC*_*i*_) explained by that eigenvector, or linear combination of parameters:

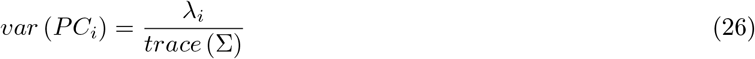

In contrast to most PCA applications where the primary focus is on principal components (PCs) with the highest eigenvalues (corresponding to directions of greatest variance) our analysis focuses on the PCs with the smallest eigenvalues. These PCs represent directions in parameter space with the narrowest posterior spread, indicating parameter combinations to which the model output is most sensitive.^47^ In other words, PCs with small eigenvalues correspond to stiff parameter combinations, while those with large eigenvalues correspond to ‘sloppy’ parameter combinations that may not be readily identifable.^47, 54^ To identify the stiff parameters and their combinations that must be constrained to achieve agreement between the model and data-derived interaction strengths, we analyze the composition of the two PCs with the smallest eigenvalues, shown in Figure 12. Parameters with strong contributions to these PCs are selected. Additionally, we perform a variance reduction analysis (not shown) that quantifies the variance changes between the initial and final SMC populations for each parameter; the results corroborate the sensitivity analysis derived from PCA.

**Figure 12.**
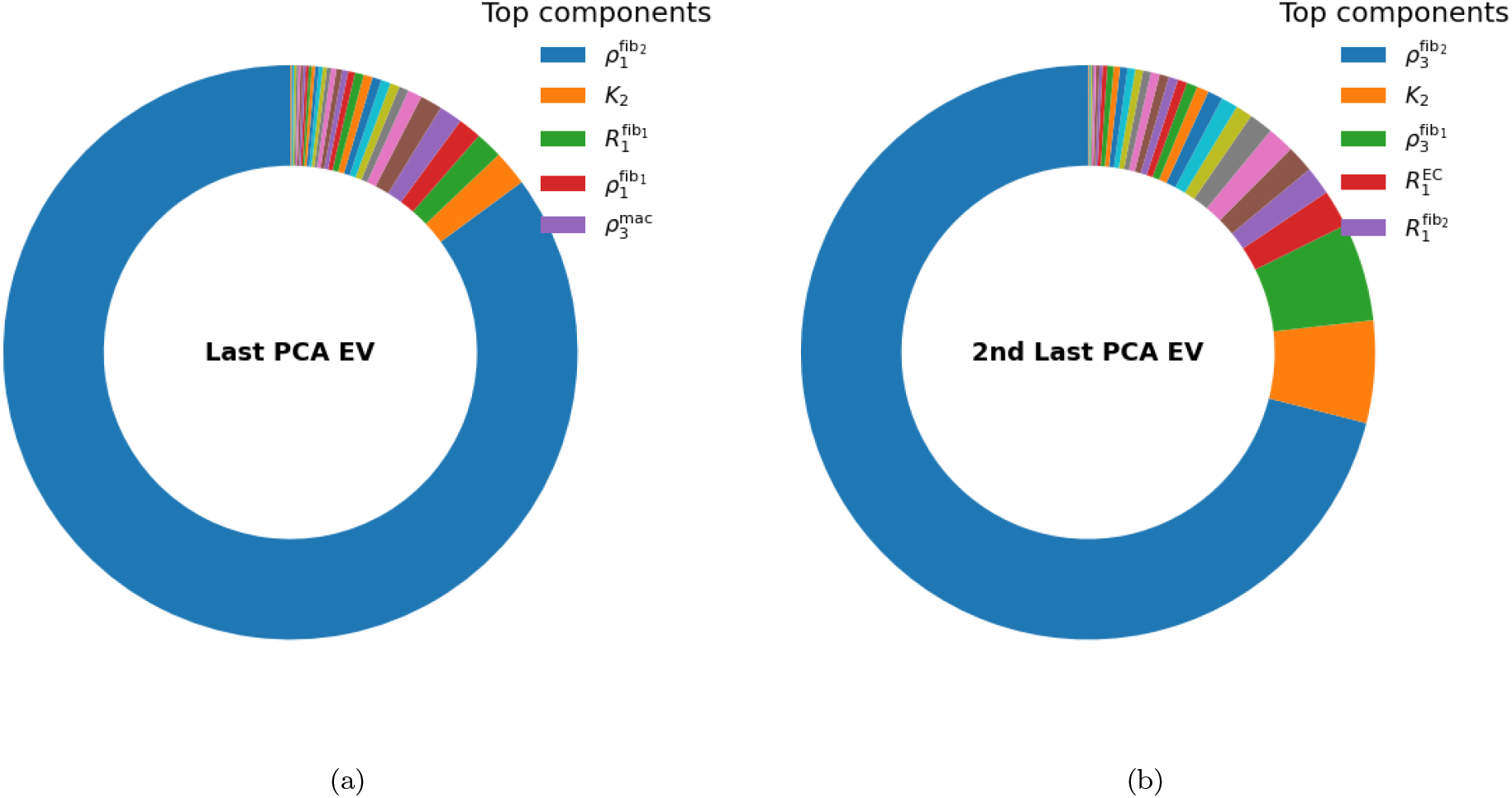
Constituent makeup of the two principal components (PCs) with the lowest eigenvalues, computed from the covariance matrix of the ensemble of particles in the final SMC population. These PCs correspond to directions of lowest variance. Parameters with prominent contributions in the standardized space indicate a high degree of constraint.

The parameters that are not prominently featured in the low-variance PCs from the SMC analysis, which likely exhibit a lower degree of constraint, are assigned to the Grade 2 category. The remaining SMC parameters, which exhibit high sensitivity and constraint according to the PCA results, are assigned to Grade 3. Together with the Grade 1 parameters from the rejection sampler, this yields three categories in total, ordered from lowest to highest constraint. These categories guide the allocation of relative learning rates for the gradient descent stage, as described below.

The final stage of the parameter calibration pipeline is gradient descent. We begin by incorporating literature-derived parameter values to define the permissible ranges used in the regularization loss function (Equation 18). We then employ the Adam optimizer, which in our experience has demonstrated fast convergence and smooth progression toward local minima. For initialization, *n* = 100 realization seeds are drawn from the particles of the last SMC population; these points are already close to minima and serve to probe the optimization landscape effectively. Each seeds is independently optimized using 3000 epochs to obtain the corresponding local minimum.

As discussed in Section 2.4.1, parameters with greater constraint should be assigned smaller learning rates. This reflects their reduced flexibility during optimization and prevents overshooting in regions where the plausible parameter space is narrow. By this stage, all parameters have been assigned to Grade 1, Grade 2, or Grade 3, in increasing order of constraint, based on the ABC rejection sampler and SMC post-hoc analyses.

This relative degree of constraint can be reflected in the learning rate *d*_*i*_ for each parameter Θ_*i*_ as:

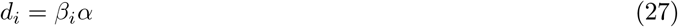

where *α* is the global learning rate and *β*_*i*_ is the parameter-specific multiplier that reflects the relative parameter constraint:

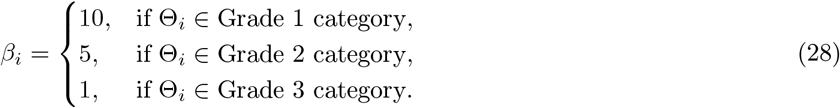

Hence, only two hyperparameters remain to be tuned: the global learning rate *α* and the regularization scaling factor *λ*. To determine their optimal values, we first generate a synthetic realization by randomly sampling from the prior distribution used in the rejection sampler step. This realization is then designated as the ‘true’ realization **Θ**^*t*^, which we aim to recover. We then compute the corresponding interaction strengths and scale them according to the bioinformatics-derived interaction strength scale (specific to the chosen tool; in this case, CellphoneDB) to produce the synthetic interaction strength dataset.

We next run the complete calibration pipeline without knowledge of **Θ**^*t*^ and select *n* = 20 seeds from the last SMC population. For each combination of *λ* ∈ {0.001, 0.005, 0.01, 0.015, 0.02} and *α* ∈ {5 × 10^*−*3^, 1 × 10^*−*2^, 2 × 10^*−*2^, 3 × 10^*−*2^}, we execute the gradient descent algorithm. The optimal hyperparameters (*λ*\*, *α*\*) are chosen as those that produce a parameter realization **Θ**^*d*^ closest to the true realization, according to the weighted relative error metric:

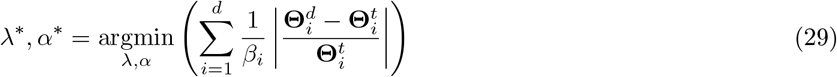

where *d* is the dimension of the parameter space, and *β*_*i*_ is defined according to Equation 28. In the case of the *TGFβ* model, the combination *λ* = 0.001 and *α* = 5 × 10^*−*3^ was found to be optimal according to Equation 29, with all constrained parameters in the Grade 2 and 3 categories falling within 10% of their respective true values. Figure 13 shows an example of the total loss across the training epochs for one of the initialization seeds, showing smooth convergence towards a low error.

**Figure 13.**
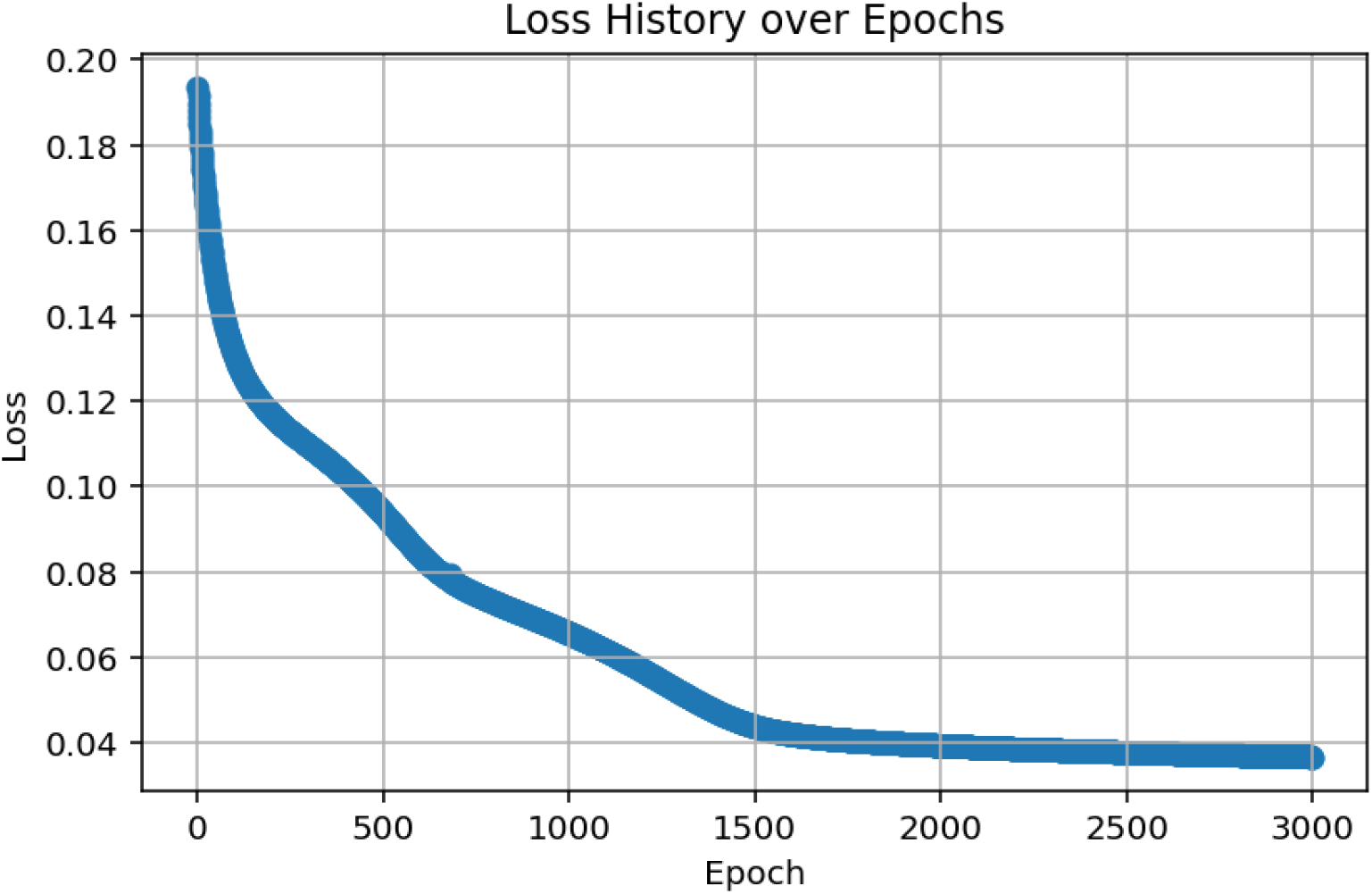
The evolution of the total loss across the epochs for one of the training seeds for the TGF*β* model.

The parameter values obtained at the final epoch of the gradient descent algorithm were used to calculate the interaction strengths between the different cell types for each TGF*β* ligand-receptor pair according to Equation 16, which were then compared against the interaction strengths derived from transcriptomics data. An example comparison using a single initialization seed is shown in Figure 14, demonstrating strong agreement between the model and transcriptomics-derived interaction strengths, with a correlation coefficient of *r* = 0.992.

**Figure 14.**
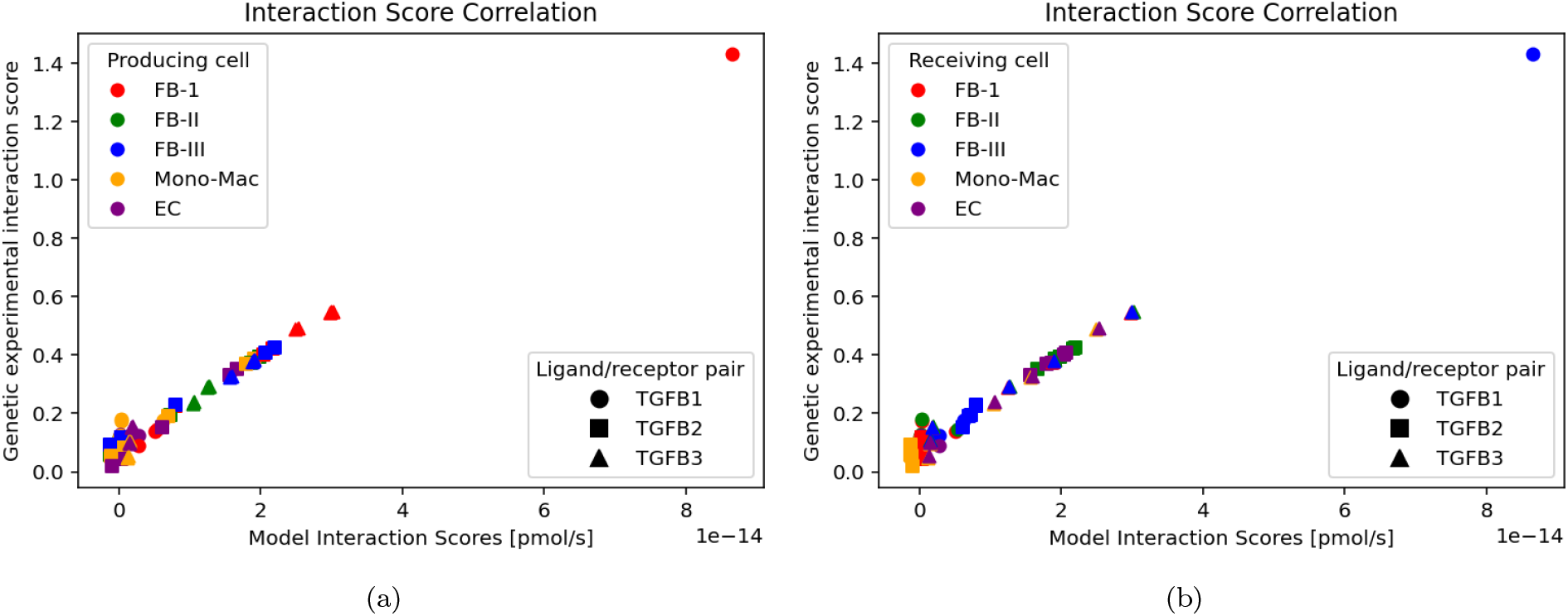
Model vs data-derived interaction strengths for the TGF*β* model for skin wound healing at day 30 post-incision, showing a strong linear relationship following parameter calibration (*r* = 0.992). The cells are colour-encoded according to TGF*β* secretion (a) and TGF*β* absorption (b). The genetic experimental interaction strengths are derived directly from the transcriptomics data using bioinformatics tools as described in Section 2.1.3 (using the CellPhoneDB cell-cell communication inference tool) or and Appendix A, while the model interaction strength scores are calculated from Equation 16.

**Figure 15.**
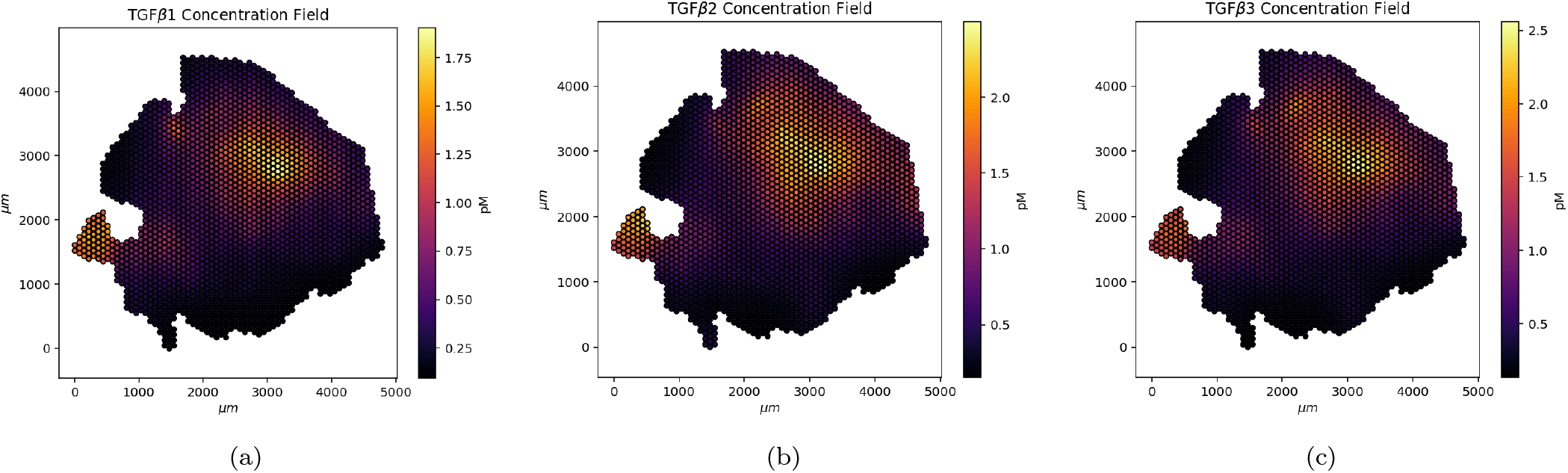
The numerically derived TGF*β* concentration fields for three ligands in the normal wound healing tissue sample at day 30 post-inicision^18^ using same parameter realization instance as in Figure 14.

These parameter realizations can also be used to numerically simulate the concentration fields of the ligands of interest by solving Equation 15 in order to examine the patterns of their spatial distribution. Figure 15 shows the TGF*β* concentration fields obtained for each ligand using the same final parameter realization instance as in Figure 14. The obtained concentration fields can also be compared with the spatial distribution of the different cell types to extract any co-localization patterns (see Section 3).

## 3 Discussion

In this paper, we have developed a computational pipeline for the parameter calibration of cellular reaction-diffusion models from transcriptomics data. In particular, we looked at scRNA-seq and ST data as a potential rich, increasingly accessible, and perhaps underutilized source of data for the calibration of the reaction-diffusion models of ligands involved in chemical cellular communication. The computational pipeline developed draws on methods from bioinformatics, machine learning, numerical analysis, and Bayesian inference, integrating existing and new computational tools and algorithms into a single computational pipeline for the robust calibration to facilitate a rigorous and robust parameter calibration process that is physiologically informed and data-driven, both of which are important considerations if mathematical models are to have any substantial research and clinical utilities in the future.

The core contribution of this paper is the hierarchical combination of the three parameter calibration schemes into a single pipeline, as illustrated in Figure 7, designed to overcome the individual limitations of each parameter calibration method. The computational pipeline developed in this study can be operated fully automatically, should the user choose. All code is freely available in an online repository. By relying solely on open-source data, the study demonstrates that the pipeline can be applied and calibrated using readily accessible information, making it especially useful for research groups without direct access to high-quality experimental datasets.

This computational pipeline is primarily intended as a proof-of-concept demonstration that genetic data can be used to parametrize higher-scale model dynamics; hence, it was designed to be flexible, allowing modifications and refinements in its assumptions, the choice of parameters, mathematical formulations, and algorithms or toolkits. Examples of potential modifications include altering the mathematical formulations describing reaction kinetics (Equations 5–7) to accommodate more comprehensive binding dynamics, different formulations for the interaction strengths in Equation 16 such that the production rate contributions could factor in ligands produced by neighbouring CVs, customized choice of the relative learning rates for different parameter categories (Equation 28), and different choice of the toolkits or algorithms for extracting interaction strengths across ligand-receptor and cell-type pairs. Below, we elaborate on these points, and highlight several potential refinements for future work.

In our main case study, the analysis is restricted to a subset of cell types: fibroblasts, macrophages, and endothelial cells, assuming that all *TGFβ* production and absorption during the remodeling phase originates from these cells. Naturally, other cell types also participate in cellular cross-talk and may contribute to abnormal wound healing. For example, Liu et al.^18^ analyzed scRNA-seq data from venous diabetic foot ulcer patients and normal wound healing patients, revealing a link between failed keratinocyte migration and impaired inflammatory responses in chronic wounds. Accordingly, the computational pipeline could be extended to incorporate the contributions of any of the other cell types identified from the scRNA-seq analysis in Section 2.1.1.

Additionally, our model relies on a quasi-steady-state approximation for *TGFβ* levels to simplify the PDE system, which is justified because during conditions such as the wound remodeling phase, TGF*β* levels and cell population dynamics stabilize and TGF*β* levels approach stable, non-zero concentrations, as described in Section 2.1.1. In certain fibrotic tissues, *TGFβ* production can be elevated and unstable due to dysregulated feedback,^55^ suggesting that the steady-state assumption may need to be revisited and the PDE formulation in Section 2.2 adjusted accordingly. Furthermore, in this study, we have relied on a point estimate of cell densities by using the 5% quantile (*q*_05_) of the cell numbers per spot and cell type across the Visium hotspots. The Bayesian inference package Cell2location, however, provides a full distribution for the cell number of each type and spot to account for uncertainty in the cell type or number obtained from deconvolution. In practice, the differences between *q*_05_ and the mean for the case studies in this paper were minimal, indicating tight distributions and low uncertainty. In future work, the computational pipeline could be extended to explicitly accommodate this uncertainty in the estimated cell numbers per type and spot.

In this paper, we have relied on a simplified kinetic representation of ligand-receptor binding and downstream signal propagation as a starting point for modeling *TGFβ* signaling in our case studies. In reality, the binding kinetics are more complex, and in certain contexts, it may be important to incorporate these complexities into the model. For example, as explained in Section 2.1.3, *TGFβ* binding involves the recruitment and formation of intermediate complexes before downstream signaling occurs. For simplicity, we have only modeled the interaction of the isoforms with their primary receptors.

Similarly, our model assumes simple reaction dynamics and does not account for receptor saturation (either primary or recruited intermediates) or competition among isoforms. It is known that multiple ligand species can bind to multiple receptor species, resulting in competition. For example, of the 1,735 (secreted) ligand–receptor pairs in the Fantom5 database35, 72% of ligands and 60% of receptors bind to multiple species.^56^ This competition between multiple molecule species is prevalent and a crucial biophysical process but it is generally ignored in cellular communication inference methods.^32^ In the particular context of TGF*β* modelling, both *TGFβ*1 and *TGFβ*3 bind to the same TGF*β* type II receptor, and all isoforms share the same intermediate protein for signal propagation. Furthermore, *TGFβ* growth factors are secreted as latent complexes that require activation before binding to receptors.^57^ This activation step was not modeled explicitly, as during steady-state conditions such as the remodeling phase, production and activation rates equilibrate. However, when steady-state assumptions no longer hold, this mechanism would need to be explicitly included via additional equations. Overall, we have chosen a simplified binding and kinetic modeling for *TGFβ* as a proof of concept. More complex and nuanced binding dynamics can be incorporated in future applications of the pipeline, depending on the specific needs of the user.

In the computational pipeline, cells are annotated by type based on the clusters identified in the scRNA-seq analysis (Section 2.1.1), enabling differentiation between subtypes within a broader parent cell type (e.g., the three fibroblast clusters shown in Figure 1(a)) according to their transcriptomic profiles. This is valuable because it allows explicit separation of cells from the same parent type that may nonetheless occupy different positions along a transition or differentiation spectrum. However, within a given identified cluster or subtype, a single run of the computational pipeline implicitly assumes that all cells share identical parameter values, such as ligand production rates or receptor numbers, which may not fully reflect biological variability. In future extensions, the algorithm could be adapted to assign parameter distributions to each cell type or subtype, allowing values to be sampled and distributed across the cells in the Visium spots within that category.

We note that the computational pipeline does not perform rigorous parameter sensitivity^58^ or structural identifiability analyses,^52^ as these are computationally demanding and difficult to incorporate into a fully automated workflow. Instead, our framework leverages an automated post-hoc analysis of the realization ensembles generated by the rejection sampler and SMC schemes. These analyses, based on KS tests, PCA, and variance reduction methods, still provides a robust and insightful assessment of parameter constraints, sensitivity, and identifiability. Importantly, they can be conducted in an automated manner without requiring additional simulations.

At this stage, the relationship between the interaction strengths derived from the ligand concentration fields of the model and those inferred from transcriptomic gene expression patterns has not been clearly established. Therefore, in this study we adopt Pearson’s correlation coefficient (*r*) as a simple, relevant, and interpretable measure of compatibility to be optimized. In the future, as this relationship is better characterized, for example, through synthetic simulations, alternative evaluation metrics may be employed.

Finally, in this study, we inferred cell interaction strengths using CellPhoneDB, which estimates interactions based on the mean ligand and receptor expression between cell type pairs. These values were then compared to the values obtained from Equation 16, which computes the interaction strengths from the ligand concentration fields as well as the parameter values, and quantifies the contribution of a particular cell type to the ligand concentration and binding within a given control volume. It should be noted, however, that CellPhoneDB does not incorporate spatial information regarding cell locations or distances between interacting cells, which is important since ligands and receptors often interact within limited maximal diffusion spatial ranges.^32, 59^ Nonetheless, even without explicitly factoring in spatial information, it has been demonstrated that CellphoneDB (along with CellChat and a few other packages) can achieve over all very good performance in terms of consistency with spatial tendency and software scalability when compared to other available tools for dissecting cell-cell communications.^60, 61^

We used CellPhoneDB in this proof-of-concept study due to its ubiquity, extensive validation, widespread adoption, and relative ease of use.^6, 23, 29, 30^ Notably, a few other new toolkits and extensions to existing ones have been very recently developed to incorporate spatial information when inferring cellular communication and interaction strengths from ST data.^32, 59, 62^ To demonstrate how our computational algorithm can accommodate the employment of other bioinformatics tools to derive the interaction strengths, we employ COMMOT (COMMunication analysis by Optimal Transport), which accounts for the competition between different ligand and receptor species as well as spatial distances between cells,^32^ to infer the cell-cell type interaction strengths from a sample of acne keloidalis^23^ in Appendix A. While existing bioinformatics tools for inferring cell–cell communication from transcriptomic data share similar underlying frameworks, each emphasizes different aspects of interaction networks, and their complementary strengths make them valuable when used together. Here, we present our pipeline primarily as a proof of concept, which can be extended or adapted by other researchers depending on their specific needs and the tools available; hence, comparative evaluations of these tools have been conducted elsewhere^60, 61^ and are beyond the scope of this study.

Our main focus is describing the development and rationale behind the algorithms presented in this study, and the way they can be deployed sequentially combined to support parameter calibration, as exemplified via the case studies on normal and abnormal wound healing (Sections 2 and Appendix A, respectively). A detailed analysis of the model results, including the final parameter values and any derivative outputs, is beyond the scope of this paper. In theory, the post-hoc analysis of the parameter values obtained from the calibration pipeline, as well as the resulting concentration fields of the ligands obtained from simulations along with their colocalization patterns with the spatial distributions of the different cell types, could provide valuable insights into the biophysical mechanisms of normal physiological processes, as well as those underlying disease progression.

For example, even before using the obtained parameter values to conduct dynamic simulations, examining the final model parameter values in the context of normal and abnormal skin wound healing could pinpoint the origins of disruptions in cell signalling, whether they arise due to altered production rates of a specific TGF*β* isoform by a particular cell type, increased receptor presentation on another cell type, or changes in how ligands interact with the extracellular environment, thereby affecting the diffusion coefficient *D*_*g*_ or the decay rate *λ*_*g*_. Furthermore, the numerically obtained concentration fields can be compared with the spatial distribution of different cell types to identify potential colocalization patterns. For example, in Figure 15, two clusters of high TGF*β* concentrations are visible, as indicated by the yellow regions in the heatmap. Similarly, Figure 3 shows two regions with high fibroblast and macrophage densities that are colocalized. The two clusters of high TGF*β* concentrations coincide with the two clusters where fibroblast and macrophage cells are colocalized, potentially indicating increased TGF*β* activity and thus cellular cross-talk between these cells.

## 4 Acknowledgements

We would like to thank Nicholas Dawes, Scientific Computing Engineer at the University of Dundee, for his guidance and support with submitting the computational runs on the Dundee HPC. This work was supported by the Agence Nationale de la Recherche (ANR) under grant number ANR-21-CE45-0025-01.

## 5 Data Availability

The transcriptomics datasets analyzed in this study are publicly available in the GEO database under accession numbers GSE241132 and GSE206790, corresponding to the original studies by Liu et al.^18^ and Hong et al.^23^, respectively. The code for the computational pipeline used for analysis will be made publicly available in a public repository (GitHub) at the time of publication.

## A Applying the Calibration Pipeline on a Acne keloidalis Tissue Sample

In this Section, we use the computational pipeline to calibrate the TGF*β* model parameters from a transcriptomics sample of a patient suffering from chronic acne kelidalis.^23^ Acne keloidalis is a primary scarring alopecia that is characterized by prolonged inflammation in the scalp, resulting in keloid-like scar formation and hair loss. Histologically, acne keloidalis is characterized by a granulomatous reaction and extensive fibrosis in the later chronic stages.^23^ Figure 16(a) shows the histological image of the tissue sample, while Figure 16(b) displays the corresponding Visium spots arranged in a hexagonal grid and overlayed on the tissue section. The annotated clusters and cell type obtained from the scRNA-seq data analysis are shown in Figure 17.

**Figure 16.**
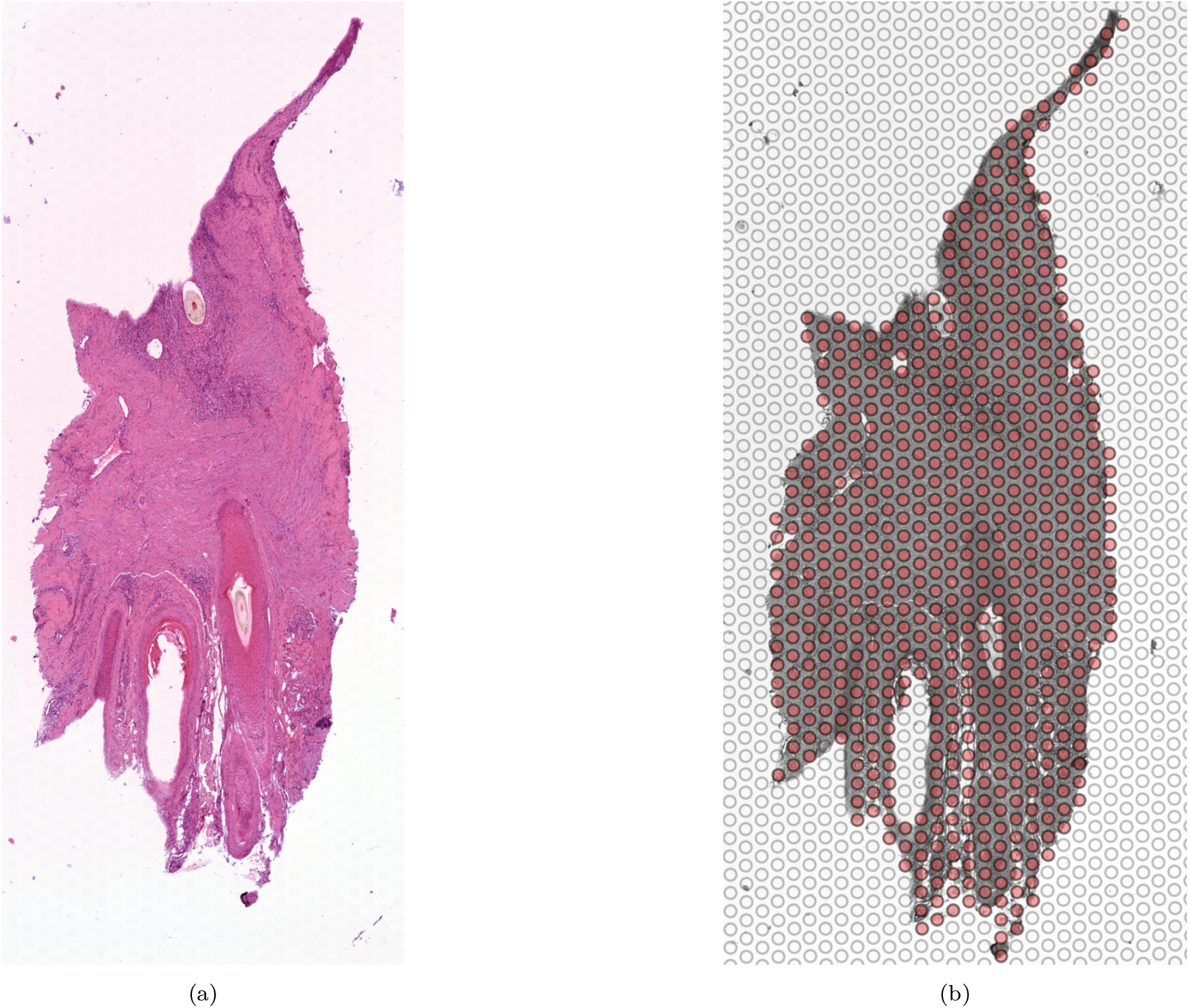
Histological tissue sample representing chronic acne keloidalis (a), with the corresponding Visium spots in red arranged in a hexagonal grid and overlaid on the tissue sample (b). Figures derived from GEO accession GSE206790, originally published by Hong et al.^23^

**Figure 17.**
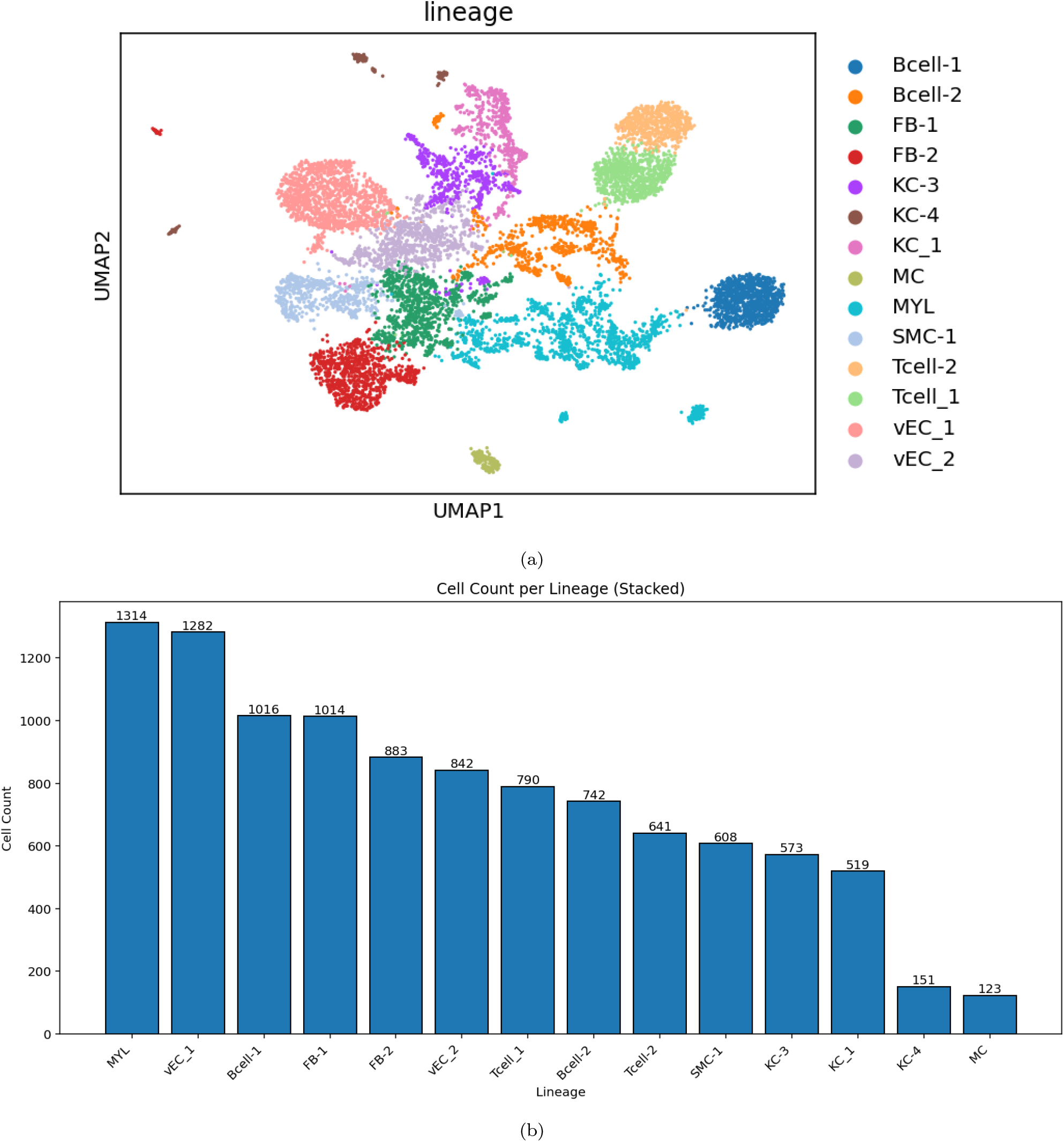
The processed raw scRNA-seq data for the acne keloidalis sample from the study by Liu et al.^23^ (a) The UMAP embedding of the identified clusters with the corresponding cell type annotations. (b) The cell count per cell type.

As discussed in Section 3, the computational pipeline accomodates the use of different bioinformatics tools to derive the interaction strengths from the transcriptomics data, depending on the nature of the sample and the needs of the user. To showcase the modularity of the pipeline, in this Section we use COMMOT (COMMunication analysis by Optimal Transport), a recently developed bioinformatics tool that can be used to infer the interaction strengths from spatial transcriptomics data.^32^ Unlike traditional approaches that rely on data from scRNA-seq, COMMOT analyses ST data and explicitly accounts for the competition among ligand-receptor pairs, as well as spatial distances between spots by introducing a constraint on the maximal distance (default = 500*µm*) across which cells could interact; the latter is particularly important, since direct cell–cell communication via ligand secretion can occur only within limited spatial ranges governed by diffusion.^32, 59^ By incorporating these constraints, COMMOT reduces the risk of false positives inherent in methods that ignore spatial context.^32^

The output of COMMOT for a given ligand–receptor pair (or sum of pairs) applied to a Visium transcriptomics dataset is a *spot* × *spot* interaction matrix, where the (*i, j*)^th^ entry represents the interaction score resulting from ligands secreted by spot *i* binding to the designated receptors on spot *j*. In our analysis, we apply a spatial distance limit of 500, *µm* for inference.^32^ To assess potential interactions, including heterodimeric complexes, COMMOT can utilize curated databases from either CellChat^59^ or CellPhoneDB,^29^ the two most widely used tools for inferring cell–cell communication networks. Figure 18 compares the inferred spatial interaction strengths, summed over all relevant direct TGF*β* interactions (TGF*β* ligands binding to TGF*β* receptors), between sending and receiving spots using the CellPhoneDB versus CellChat databases, demonstrating overall general agreement. For subsequent analyses, results obtained using the CellPhoneDB database are employed.

**Figure 18.**
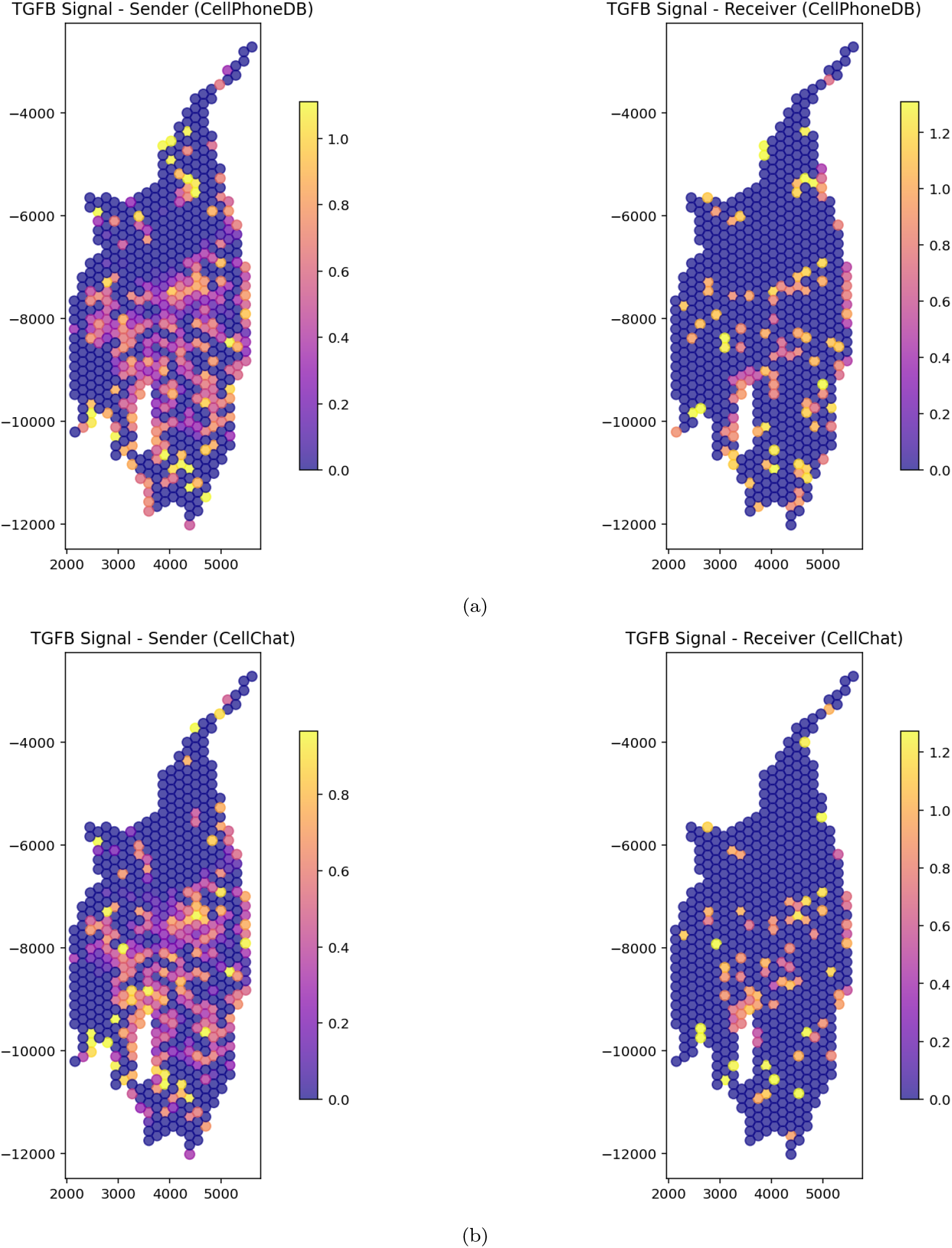
The spatially inferred spot-spot interaction strengths obtained from using COMMOT on the ST sample transcriptomics data for acne keloidalis,^23^, when using the CellphoneDB (a) and CellChat (b) ligand-receptor curated databases.

**Figure 19.**
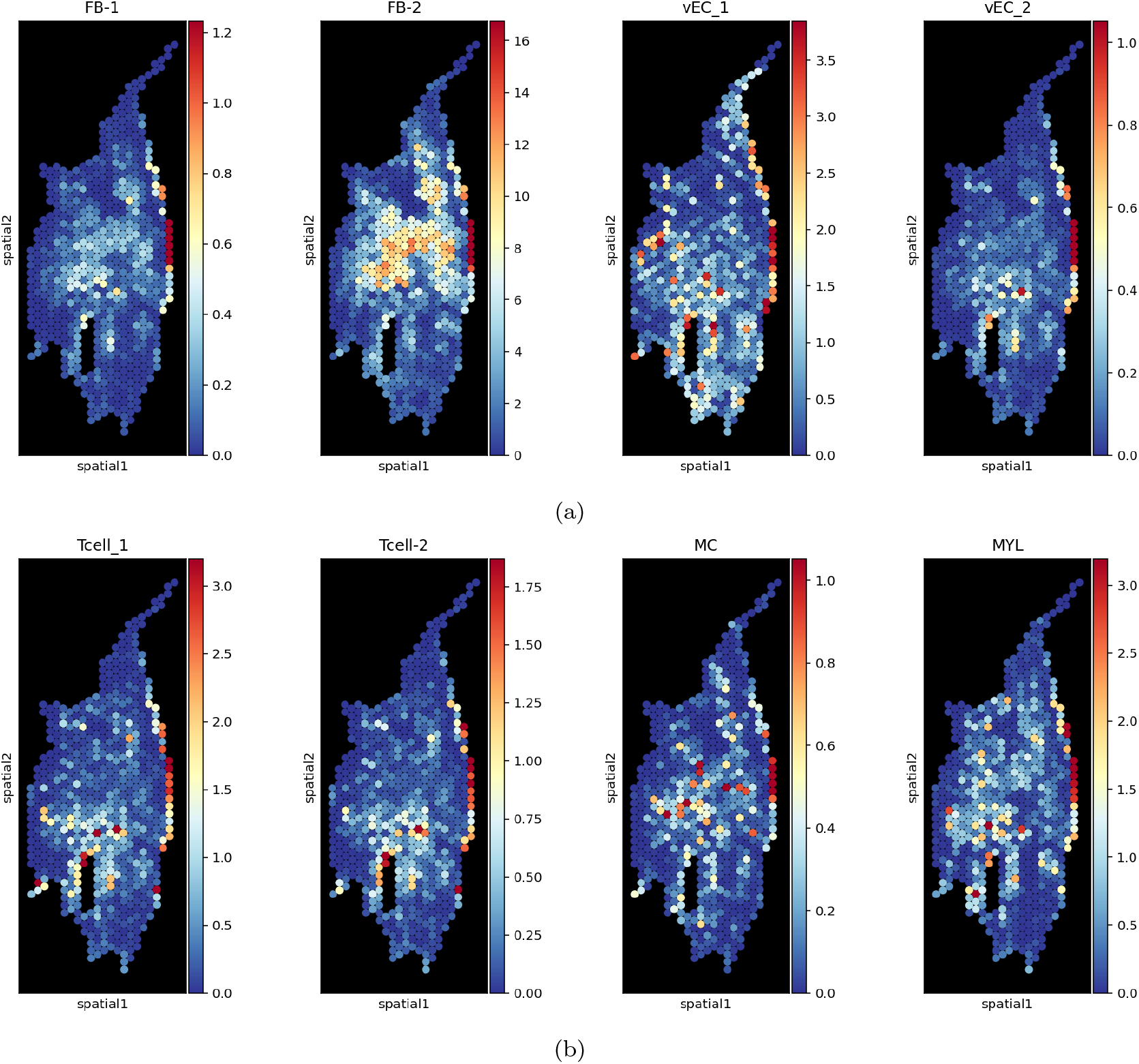
he spatial distribution across the Visium spots of the fibroblast and endothelial cell subtypes (a) and the immune cell subtypes (b) in the acne keloidalis sample.^23^ The cell counts represent the 5% quantile of the posterior distribution obtained from the Cell2location analysis. FB: fibroblast, vEC: vascular endothelial cells. MC: Macrophages. MYL: Myeloid cells.

The next process in the computational pipeline is to approximate the cell numbers per spot and cell type. Again, the Cell2location toolkit is used and the *q*_05_ estimates for the fibroblasts, endothelial and immune cell subtypes are obtained, shown in Figure 19.

The spatially inferred interaction strengths from COMMOT, however, are only between the Visium spots and do not incorporate any knowledge of the cell types, their abundances, annotations, and gene expression profiles, features typically obtained from scRNA-seq analyses. In contrast, our computational pipeline requires interaction strengths to be specified at the cell type level, in the form of a *cell type × cell type* matrix *M*_*AB*_ for each ligand–receptor pair (*A, B*), where rows and columns represent the receiving and producing cell types, respectively. By comparison, COMMOT outputs a *spot × spot* matrix *S*_*AB*_, which must be transformed to align with our framework.

To transform *S*_*AB*_ into the cell type–level matrix *M*_*AB*_, we leverage the proportions of each cell type within each spot which can be derived from the Cell2location analysis, along with the mean ligand and receptor expression levels in each cell type from the annotated scRNA-seq dataset, as follows:

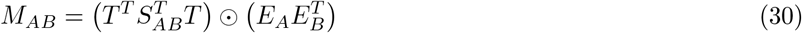

Here, *T* is the *spot* × *cell type* matrix of cell type proportions per spot, derived from Cell2location’s cell number estimates. *E*_*A*_ and *E*_*B*_ are vectors of length equal to the number of cell types, containing the log-normalized mean expression of ligand *A* and receptor *B*, respectively. The symbol ⊙ denotes element-wise (Hadamard) multiplication. After obtaining the *cell type* × *cell type* interaction matrix *M*_*AB*_, the subsequent analysis can proceed analogously to the framework described in Section 2.

## B Particle-based Model Mathematical Derivation

For particle-based models where the source and sink terms are localized to point sources, *S*(*x*) can be formulated using the Dirac delta function *δ*:

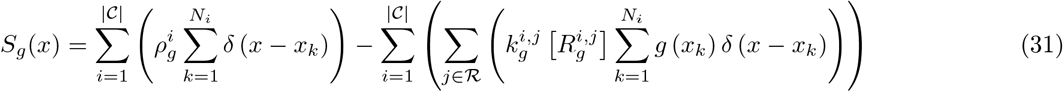

where |𝒞| is the cardinality of 𝒞, or the number of cell types, *N*_*i*_ is the number of cells belonging to to cell type *i*, and *j* ∈ ℛ refers to the receptor(s) binding ligand *g*. Given that the source and sink terms are represented by point sources, and that inserting Equation 31 into Equation 8 results in a linear spatial PDE, the global solution for *g*(*x*) can be analytically formulated via the superposition of the Green functions:

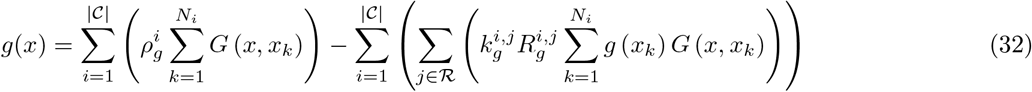

For the steady-state (time-independent) 2D case, the Green function *G*(*x, x*_*k*_) (*s/m*^2^) can be expressed in terms of *K*_0_, the zeroth-order modified Bessel function of the second type:

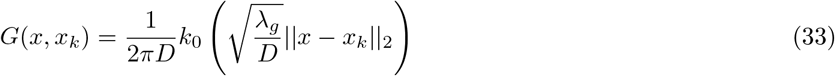

Rearranging Equation 32, we can solve for the growth factor concentration at any cell with location *x*_*k*_:

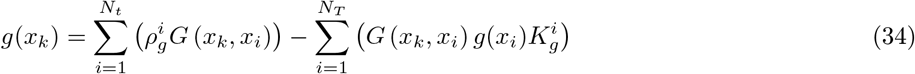

where, for the sake of mathematical convenience, *N*_*T*_ corresponds to the total number of cells (of types of interest) in the domain, and 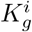 is the ‘equivalent’ binding rate for growth factor *g* on cell *i*, formed from the summation of the binding rates of the different receptors for *g* on cell *i*.

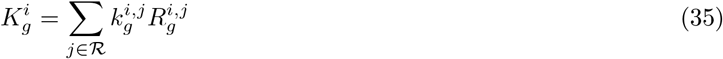

Just as for the continuous formulation, Equation 34 can be arranged to form a linear system of Equations to solve for **g**_**k**_, the vector corresponding to growth factor concentration at all cell centres:

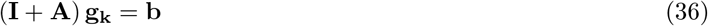

where the entries for **b** and **A** are given by:

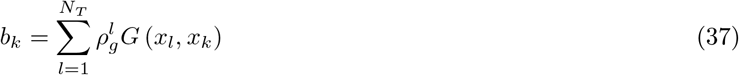

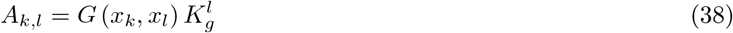

It’s important to note that the Green’s function solutions, resulting from the assumption of highly localized Dirac delta source/sink points, exhibit both numerical and physical singularities at those points. These singularities, in turn, influence the overall solution obtained from the superposition of the Green functions. This can be seen numerically when evaluating *G*(*x*_*i*_, *x*_*k*_) for the case when *x*_*i*_ = *x*_*k*_: for small distances ||*x*_*i*_ *x*_*k*_||_2_ *<< l*, the second type modified Bessel function of zeroth order can be approximated as:

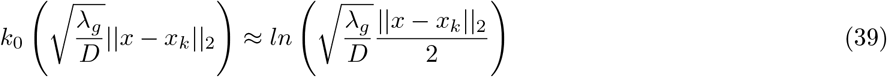

which gives rise to a singularity when ||*x*_*i*_ *− x*_*k*_||_2_ = 0. To circumvent such singularities, a simple remedy can be to calculate 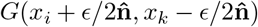 for a small constant *ϵ* and unit normal vector 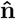 pointing from *x*_*i*_ to *x*_*k*_, such that:

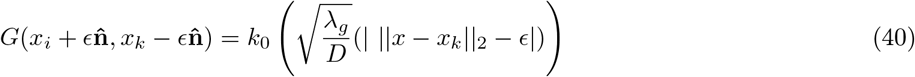

In physical terms, if *ϵ* represents the typical cell dimensions, such as the average diameter of the cells of interest, replacing Equation 33 with 40 can be viewed as assessing the impact of ligand production and absorption by ligand receptors on the cell surfaces, rather than at the cell centres. This approach is more physiologically relevant, as real systems, like this one, involve sources and sinks (e.g., cells) with finite sizes. It’s important to note that this regularization or offset still treats cells as point sources and sinks for simplicity, rather than as entities with surfaces where ligand binding and production take place. Nevertheless, this offset can provide a more accurate representation of ligand diffusion and receptor binding, while also helping to maintain the numerical stability and smoothness of the solution.

Next, for the particle-based approach, we can approximately isolate the total contribution of cell type *l* to ligand *A* binding to receptor *B* on cell-type *r* as:

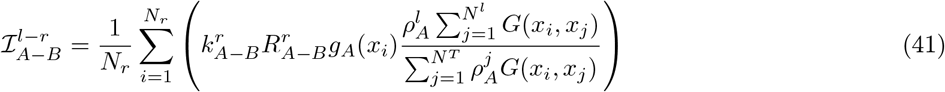

where *N*_*r*_ is the number of the receiving cells, *g*_*A*_(*x*_*i*_), *i* = 1…*N*_*r*_ are the concentrations of the growth factor *A* at the location of the receiving cells (where it binds to the receptor), 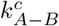 is the binding rate between ligand *A* and receptor *B* on cell type *c, N*^*l*^ is the number of cells of type *l*, and *N*^*T*^ is the total number of cells.

Since the ligand concentration field can be expressed as a linear system of equations (Equation 36), and the interaction strengths for a given realization can be determined from this field and a subset of the model parameters (Equation 41), the subsequent parameter calibration of the particle-based model can proceed analogously to the continuum-based approach described in Section 2.4.

